# Language usage modulates the neural mechanisms of selective attention in bilinguals

**DOI:** 10.1101/2025.11.26.690745

**Authors:** Stephen Theron-Grimaldi, Julia Schwarz, Pepita Alex, Mirjana Bozic

**Affiliations:** Department of Psychology, University of Cambridge, Downing Street, Cambridge CB1 3EB; Basque Center on Cognition, Brain and Language - Mikeletegi Pasealekua 69, Donostia / San Sebastián 20009; School of Psychology, University of Aberdeen, King’s College, Aberdeen AB24 3FX

**Author notes:** Corresponding author Stephen Theron-Grimaldi Department of Psychology University of Cambridge Downing Street, Cambridge CB2 3EB.

**Keywords:** bilingualism, selective attention, language usage, alpha oscillations, speech tracking

## Abstract

The use of multiple languages modulates the neural mechanisms of selective attention, but it is unclear whether these adaptations require continuous engagement and active second language usage. Here we examined whether language usage shapes selective attention in bilingualism, and what neural processes might this engage. 48 highly proficient English-French bilinguals listened to naturalistic speech streams in their first or second language, paired with either linguistic and non-linguistic interference. Participants were matched on their L2 proficiency, but were either Active, Moderate, or Inactive users of their second language. The results revealed usage-related modulation of oscillatory activity in the alpha band, with more efficient inhibitory control in Active and Moderate users leading to behavioural resilience to interference. In contrast, usage did not affect lower-level perceptual tracking of speech, as captured by mTRF decoding of speech envelopes across frequency bands. Taken together, our findings show that resilience to interference during language processing is not dependent on the perceptual speech tracking, but rather on the capacity to recruit higher-level attentional control mechanisms, a process that is dynamically shaped by bilinguals’ L2 usage.

## 1. Introduction

Bilingualism, the capacity to understand and speak more than one language, is increasingly recognised not only as a valuable skill, but also as a lens into the brain’s remarkable ability to adapt and reorganise (Li et al., 2014; Tao et al, 2021). Comparisons between monolinguals and bilinguals offer strong evidence of neural adaptation in the bilingual brain, from local structural differences to large-scale functional reorganisation (De Luca et al., 2019; Luk et al., 2011; Sulpizio et al., 2020). However, it is unclear to what extent these adaptations change dynamically with active language use. This study investigates how neurocognitive adaptation of selective attention in bilinguals might be shaped by the degree of usage of the second language.

### 1.1 The variables driving bilingualism adaptation

There is abundant evidence that bilingualism shapes both neural structure and function. For example, MRI studies have reported structural differences in the grey-matter volume of both language-specific (Olsen et al., 2015) and domain-general regions (Abutalebi et al., 2015; Mechelli et al., 2004). White-matter adaptations have likewise been documented, including greater frontal white-matter volume (Olsen et al., 2015) and enhanced microstructural integrity in key long-range tracts such as the corpus callosum (Luk et al., 2011) and the uncinate fasciculus (Pliatsikas et al., 2015). Functional studies offer additional evidence of bilingualism-induced neuroplasticity: compared to monolinguals, bilinguals show greater functional connectivity within language-related regions (Gracia-Tabuenca et al., 2024) and across broader executive control networks (DeLuca et al., 2020; Sulpizio et al., 2020). These adaptations also affect attentional mechanisms in bilinguals (Olguín et al., 2019; Phelps et al., 2022) and modulate neural oscillatory patterns linked to attentional control during non-linguistic tasks (Chung-Fat-Yim et al., 2023). Neural adaptations to bilingualism appear to serve complementary functions: structural changes likely reflect increased local processing demands in response to the presence of two competing languages, while functional reorganisation may enable more efficient communication across distributed control networks (DeLuca et al., 2019; Sulpizio et al., 2020). Altogether, they suggest a system-wide neural adaptation that enables bilinguals to manage multiple languages in real time, balancing activation and suppression demands in accordance with communicative goals (Green & Abutalebi, 2013). However, although comparisons between bilinguals and monolinguals have revealed important structural and functional neurocognitive differences, binary contrasts offer limited insight into the specific language experiences that give rise to and modulate such adaptations. This is because bilingualism is not a unitary condition, but a highly heterogeneous experience: individuals differ widely in when they acquire their second language, how proficient they are, and how frequently they use it; as well as in the environment in which they use their second language. To fully understand how bilingualism shapes the brain, it is therefore essential to move beyond the broad and static group comparisons and unpack the distinct contributions of these underlying factors.

One commonly studied modulator of bilingualism is the Age of Acquisition (AoA) of a second language. Several studies have shown that bilinguals who acquired their second language (L2) at an early age display reduced signatures of neural effort during language production and more coordinated recruitment of control regions, often accompanied by greater bilateral engagement of the language networks (Anderson et al., 2024; Berken et al., 2016; Gullifer et al., 2018). However, results are mixed when it comes to late bilinguals. Some studies suggest they still show distinct neural adaptations compared to monolinguals (Sulpizio et al., 2020), while others find no such effect (Gracia-Tabuenca et al., 2024). Furthermore, although AoA may modulate bilingual adaptation, it cannot capture the dynamic, experience-dependent plasticity that characterises the bilingual brain throughout the lifespan (Schlegel et al., 2012; Tu et al., 2015; Wei et al., 2024).

Another factor that has received considerable attention is language proficiency, which captures the depth and complexity of a bilingual’s knowledge of their second language. Highly proficient bilinguals have been associated with greater grey matter volume in language-related areas (Abutalebi et al., 2015; Mechelli et al., 2004), stronger functional connectivity within and between language and control networks (Sulpizio et al., 2020) and reduced cognitive effort during language control tasks (Mouthon et al., 2020). Importantly, proficiency has also been linked to brain changes in longitudinal studies, making it a useful marker of neural adaptations, with intensive language learning leading to rapid structural and functional adaptations (Mårtensson et al., 2012; Schlegel et al., 2012; Stein et al., 2012). However, proficiency is often treated as an isolated predictor, without considering its relations with language usage. As a result, it remains unclear whether observed neural changes are due to the cumulative acquisition of language skills or recent and active engagement with the language. Proficiency is also arguably relatively stable once a language has been acquired and does not capture the frequent fluctuations of bilingual experience that may influence neuroplasticity.

Among the other significant modulating factors, language usage has emerged as the most dynamic and comprehensive predictor of bilingual neuroplasticity. A growing body of evidence shows that usage modulates a wide range of neural systems, particularly those involved in cognitive control. For example, resting-state studies have demonstrated that language usage correlates with functional connectivity in control-related networks (Gullifer et al., 2018), while task-based studies have shown that bilinguals with higher language usage engage executive regions more efficiently during demanding language tasks (DeLuca et al., 2020). Crucially, active language usage has been shown to drive neural changes even over short timescales, with only one month of decreased language usage sufficient to result in measurable changes in functional brain organisation (Tu et al., 2015). Similarly, a break in language practice was associated with partial reversal of training-induced structural changes (Hosoda et al. 2013). Other longitudinal research has confirmed that brief periods of L2 training or immersion can reshape both structural connectivity and activation patterns (Hosoda et al., 2013; Pliatsikas et al., 2017; Schlegel et al., 2012; Wei et al., 2024). Beyond short-term effects, usage also appears to play a key role in the long-term maintenance of bilingual adaptation: lifelong bilinguals who continue to use their second language show preserved grey matter volume in both language-specific and control-related regions (Abutalebi et al., 2015). Higher levels of language usage have also been associated with increased white matter volume (Olsen et al., 2015) and greater white matter integrity (Luk et al., 2011) and may provide a protective effect against age-related decline (Perani & Abutalebi, 2015). These findings are well captured by the Dynamic Restructuring Model (Pliatsikas, 2020), which emphasises the acquisition and maintenance nature of neuroplasticity in bilingualism.

### 1.2 Selective Attention as a Usage-Dependent Mechanism

Cognitive control is a domain in which usage-dependent plasticity may be particularly important. Bilinguals often outperform monolinguals on tasks that rely on inhibition, conflict monitoring and flexible shifting, including the Flanker and Simon paradigms (Filippi et al., 2012; Luk et al., 2010) as well as syntactic ambiguity resolution (Filippi et al., 2015). Converging neural evidence supports these behavioural findings, highlighting a more efficient control system in bilinguals. This is reflected in earlier attentional ERP responses (Kuipers & Thierry, 2015), reduced anterior cingulate activation during conflict resolution (Liu et al., 2021), and enhanced functional integration at rest (Gracia-Tabuenca et al., 2024). The Adaptive Control Hypothesis (Green & Abutalebi, 2013) attributes such effects to the continual need in bilinguals to select the appropriate language while inhibiting the non-target one. Crucially, it suggests that the strength of such control adaptations may vary with ongoing language engagement. A small but growing body of research supports this view, with several studies showing that variation in everyday language use predicts differences in executive performance. For instance, Hartanto and Yang (2016) found that frequent code-switchers exhibited smaller task-switching costs, and Kheder and Kaan (2021) showed that higher daily L2 use predicted reduced Simon interference. These effects were observed even when controlling for age of acquisition (AoA) and proficiency, suggesting that usage plays a distinct role in shaping cortical control networks.

In the domain of selective attention specifically, the available evidence shows robust effects of bilingualism on underlying neurocognitive processes. In both speech-in-noise and dichotic listening paradigms, bilinguals tend to show enhanced stability of sound encoding (Krizman et al. 2012; 2014; 2021), as well as different patterns of neural activations compared to monolinguals when focusing on auditory targets in the presence of interference (Olguin et al, 2019; Phelps et al, 2022). In such cases, monolinguals typically show robust effects of the intelligibility of interference on the encoding of attended speech (Olguín et al., 2018), sometimes accompanied by corresponding behavioural costs (Filippi et al., 2012, but see Olguin et al, 2019). Bilinguals tend to maintain stable comprehension (Filippi et al., 2015; Olguin et al, 2019; Phelps et al., 2022) and preserve more consistent cortical tracking of the attended stream irrespective of the type of interference (Olguín et al., 2019). These findings suggest that bilinguals develop a selective attention adaptation, likely supported by the habitual need to attend to one language while inhibiting another, which may allow them to filter out effectively both intelligible and unintelligible distractors.

However, despite the existing evidence about the modulation of selective attention in bilingualism, some fundamental questions remain open. First, while theoretical accounts such as the Adaptive Control Hypothesis (Green & Abutalebi, 2013) suggest that language-driven cortical adaptations develop through bilingual experience, it is unclear whether such changes to the mechanisms of selective attention rely on active L2 usage. If the ability to effectively resist disruption from interference arises from the repeated need to select the target language and suppress the non-target one, as postulated by the Adaptive Control Hypothesis (Green & Abutalebi, 2013), then usage can indeed be expected to play a central role in bilingual adaptation of selective attention. As a result, greater current L2 usage is expected to lead to more robust adaptation, while this might diminish or even disappear when the second language is no longer used. A related question is therefore how much usage is needed to sustain these attentional adaptations? While prior research suggests that neural adaptations may decline following reduced language usage (Hosoda et al., 2013; Tu et al., 2015), recent findings highlight non-linear effects, with certain adaptations plateauing after a given threshold of experience (Korenar et al., 2023).

Secondly, it is not clear what neurocognitive mechanisms might underpin the effects of usage on selective attention in bilinguals. Much of the previous research on neural adaptation of selective attention in bilinguals focused on cortical tracking of the speech signal, quantifying how closely the EEG activity encodes the acoustic signal present in the speech envelope (Blanco-Elorrieta et al., 2020; Olguin et al, 2019; Phelps et al, 2022; see Rossi et al. 2023 for review). Envelope tracking is a well-established index of attentional processing, reflecting the findings that the temporal envelope of speech is both strongly represented in the brain (Crosse et al., 2016; Lalor & Foxe, 2010) and robustly influenced by selective attention (Ding & Simon, 2013; Fuglsang et al., 2017; O’Sullivan et al., 2015). Applying this paradigm to attentional processing in bilinguals revealed that their cortical tracking of attended speech is less susceptible to interference manipulations compared to their monolingual counterparts (Olguin et al, 2019; Phelps et al, 2022). Yet in addition to broadband tracking of the speech envelope, cortical tracking in specific frequency bands was shown to provide valuable further evidence about the nature and the source of the observed effects. For instance, cortical tracking of speech in the delta band (1-4Hz) was shown to relate to higher order, top-down syntactic and semantic integration and processing of sentential structure, as well as effortful speech processing in difficult listening conditions (Ershaid et al, 2024; Ding et al., 2016; Lu et al, 2023; Mai & Wang, 2019; Molinaro & Lizarazu, 2018). Speech tracking in the theta band (4-8Hz), in contrast, is thought to align with the syllabic rhythm of speech and is thus considered sensitive to bottom-up acoustic fluctuations and regularities in the signal (Di Liberto et al., 2015; Doelling et al., 2014; Mai et al., 2016; Peelle et al., 2013). Comparing the tracking in these two frequency bands in bilinguals could therefore shed light on the source of the possible usage effects, and whether this may be affecting top-down integration or more low-level, bottom-up processing of the attended speech signal.

A complementary mechanism underpinning selective attention to spoken signal is power modulation across speech- and attention-relevant frequency bands (Tune et al., 2021). Recent work has shown that speech intelligibility affects cortical encoding and power modulation differently (Hauswald et al., 2022; Mai & Wang, 2023), and that examining these two neural markers together provides a more comprehensive picture of attentional dynamics and listening success (Drgas et al., 2021; Tune et al., 2021). A particular focus here is on the alpha band power (8-12Hz), which is thought to reflect neural filtering mechanisms that support selective attention during listening (Klimesch, 2012; Tune et al., 2021). Alpha power typically increases when irrelevant inputs must be actively inhibited (Benedek et al., 2014; Deng et al., 2020; Strauß et al., 2014; Wöstmann et al., 2017) but can decrease when the task becomes too demanding due to attentional overload (Krause et al., 2000; Obleser et al., 2012; Wisniewski et al., 2017). Research on bilinguals shows that alpha power is sensitive to AoA and increases when listening to L2 compared to L1; with both of these effects indicative of additional attentional control (Grant et al., 2022). Jointly, cortical speech tracking and power analyses can thus provide complementary information on the attentional mechanisms in bilinguals, potentially allowing us to pinpoint more precisely the likely source of any usage effects on their attentional processing.

### 1.3 Current study

We aim to answer two central questions here. First, to what extent does language usage shape the adaptation of selective attention in bilingualism? While previous research established that bilingualism modulates the neural mechanisms of selective attention, it remains unclear whether these changes are stable outcomes of past experience or dynamic adaptations that require continuous engagement with multiple languages. To address this, we tested highly proficient English-French bilinguals who were matched in their L2 proficiency and learning history, but varied in the degree of their current L2 usage. The were divided into three groups: those who used their L2 daily (active users); those who used it only occasionally (moderate users); and those who used their L2 only very sporadically (inactive users). Based on the evidence reviewed above, we expected that active or even moderate L2 usage should lead to more prominent effects (Green & Abutalebi, 2013; Korenar et al., 2023), while disengagement with second language usage may lead to less robust attentional adaptation in bilinguals. Participants were asked to perform a dichotic listening task that manipulated both the language to be attended (L1 or L2) and the nature of the interference (intelligible vs non-linguistic). This is a naturalistic listening task that allows us to tap into the bilinguals’ processing of continuous speech under selective attention conditions. By testing selective attention to both L1 and L2 under comparable interference conditions, we additionally aimed to determine whether these adaptations are language-independent and apply across bilinguals’ languages. We predicted that any attentional effects would be consistent across L1 and L2, reflecting a general bilingual attentional mechanism rather than a language-specific process. Including both linguistic (intelligible) and non-linguistic (unintelligible) interference allowed us to test whether any usage effects may vary as a function of intelligibility of the distractor.

Our second core question was: what neural processes and mechanisms might underpin the effects of L2 usage on selective attention in bilinguals? To address this question we use two complementary analyses, cortical tracking of the speech envelope and power spectral density analysis. Multivariate Temporal Response Function (mTRF, Crosse et al., 2016, 2021) approach was used to quantify how the cortical activity tracks attended and unattended speech envelopes in the three usage groups across frequency ranges. We also examined spectral power changes across frequencies, with a particular focus on alpha oscillations (8–12 Hz) that tap into the allocation of key attentional resources. Based on previous findings, we assumed that the bilingualism-induced modulation of selective attention most likely reflects redistribution of existing processing resources commensurate to load and task demands rather than a fundamental change to how the speech signal is encoded (Phelps et al, 2022; Phelps & Bozic 2025. We therefore hypothesised that L2 usage would be less likely to affect low-level cortical tracking of speech acoustics (as captured by theta band tracking in particular), and more likely to affect the top-down mechanisms of attentional allocation required for distribution of the existing processing resources under challenging listening conditions, primarily reflected in alpha power changes.

## 2. Materials and Methods

### 2.1 Participants

Fifty-three young adults participated in the experiment. They were all native speakers of English (L1) and fully proficient in French (L2); with some also occasionally using additional languages. Five participants were excluded from the data analysis: two due to low comprehension accuracy (2σ below the average) and three due to lower L2 proficiency compared to the rest of the sample (see Section 2.2 for details). The final sample size was 48 participants (mean age: 22.3; 14 males). All participants were between 18-32 years old without any hearing, attention or language disorders. All reported acquiring English before French. The study was reviewed by the Psychology Research Ethics Committee of the University of Cambridge. Participants were compensated for their participation.

### 2.2 Demographic questionnaire and L2 proficiency

All participants completed a demographic and language background questionnaire, as well as a French proficiency test. The demographic and language background questionnaire was adapted from the Bilingual Language Profile questionnaire (Birdsong et al., 2012) and the Language and Social Background Questionnaire (Anderson et al., 2018). It assessed participants’ demographic information and language profiles, including the frequency, intensity, and modality of language usage. The French proficiency test consisted of three 2-minute audio recordings and 15 multiple-choice questions. The materials were drawn from the highest level of the DELF (Elder, 2018), a French language qualification awarded by the French Ministry of Education. While the official pass level for this test is 50%, only participants who answered more than two-thirds of the questions correctly were included to ensure an advanced level of proficiency.

Based on the language background questionnaire, participants were allocated to one of three groups depending on their frequency of L2 usage: Active-Usage (daily), Moderate-Usage (several days a week) and Inactive-Usage (fewer than once a week). Usage was assessed with a questionnaire and then verified in person. Details of bilinguals’ characteristics across Usage groups are reported in Table 1. Demographic factors (Sex, Age, Education level) and language experience (AoA, Proficiency and number of Languages Spoken) were matched between Usage groups.

**Table 1.** Bilinguals’ profile across usage groups. SD in brackets

| Characteristics | Usage |  |  | Group comparison<sup>a</sup> |
| --- | --- | --- | --- | --- |
|  | Active-<br>*N* = 16 | Moderate-<br>*N* = 16 | Inactive-<br>*N* = 16 |  |
| Demographic |  |  |  |  |
| Sex (% Male) | 38% | 31% | 19% | p = .501 |
| Age | 20.6 (2.3) | 22.3 (3.5) | 23.4 (4.0) | p = .088 |
| Education level<sup>b</sup> | 1.5 (0.8) | 1.9 (0.9) | 2.3 (1.1) | p = .093 |
| Language |  |  |  |  |
| L2 Usage<sup>c</sup> | 33.4% | 12.1% | 2.2% | p < .001 |
| L2 Proficiency score | 92.1 (9.1) | 91.2 (9.0) | 87.1 (7.9) | p = .178 |
| L2 AoA | 4.4 (3.9) | 6.2 (4.0) | 5.2 (3.7) | p = .356 |
| Languages spoken<sup>d</sup> | 2.8 (0.7) | 2.3 (0.5) | 2.4 (0.5) | p = .108 |

<sup>a</sup> Kruskal-Wallis test; <sup>b</sup> 1 = A-Levels, 2 = Undergrad, 3 = Masters, 4 = PhD; <sup>c</sup> Reported proportion of time spent using French and other non-native languages (French was dominant in all cases); <sup>d</sup> Total number of languages spoken, including English (L1) and French (L2)

### 2.3 Experiment design and stimuli

The experiment used a dichotic listening paradigm in which participants were presented with two simultaneous audio streams and instructed to attend to one while ignoring the other (Figure 1A). The attended stream consisted of 8 children’s stories in participants’ L1 (English, 4 stories) or L2 (French, 4 stories). The unattended stream consisted of 4 different children’s stories in the same language as the attended stream and 4 acoustically matched non-linguistic signals called Musical Rain (MuR). MuR was derived from unattended stories by extracting their speech envelopes and filling them with 10ms fragments of synthesized speech jittered in frequency and periodicity (Bozic et al. 2010), which preserves low-level speech features while removing intelligibility. This created four experimental conditions, in which participants attended to a narrative in either their L1 or L2, paired with either non-linguistic (unintelligible) or linguistic (intelligible) interference in the same language (English-MuR, English-English, French-MuR, and French-French, Table 2).

**Figure 1.**
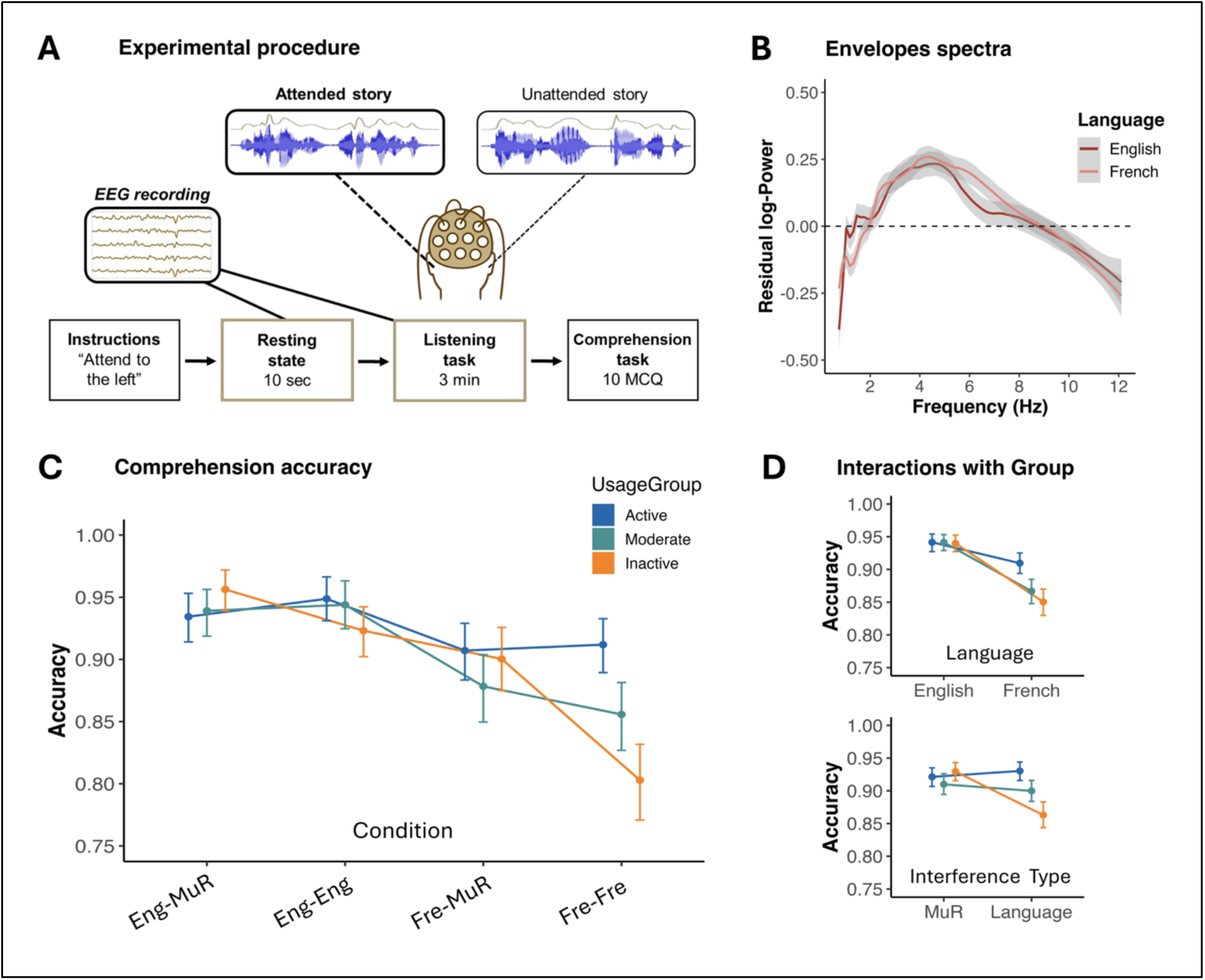
Overview of experimental design, stimuli, and behavioural results. **(A) Experimental procedure**, showing that participants listened to two concurrent audio streams (attended vs unattended) and answered comprehension questions after each block. **(B) Envelope spectra** of English and French speech stimuli, showing matched spectral profiles across 1–12 Hz with a dominant peak around 4–5 Hz. **(C) Comprehension accuracy** across all 4 conditions (English-MuR, English-English, French-MuR, French-French). **(D) Interactions of Language and Interference with Group**, with comprehension accuracy plotted by attended language and interference type, showing overall better performance in English (L1) over French (L2), and reduced performance under linguistic interference in the Inactive group. Error bars represent ± 95% CI.

**Table 2.** Experimental conditions

| Condition | Attended | Unattended |
| --- | --- | --- |
| English-MuR | English story | Musical Rain |
| English-English | English story | Different English story |
| French-MuR | French story | Musical Rain |
| French-French | French story | Different French story |

Each story consisted of 120 sentences of similar length that were split into two blocks. Each condition included two attended stories paired with two unattended stories (or corresponding MuR), resulting in four blocks per condition, and a total of 16 blocks in the experiment. To minimise speaker-related differences (Brungart & Simpson, 2007), stories were recorded by four female native speakers: two native English speakers for the attended and unattended English stories respectively, and two native French speakers for the attended and unattended French stories respectively. Sentences were edited using Audacity (version 3.2.3, 2022) to last approximately 2.75 seconds, and a 275 ms silence gap was inserted between each sentence before concatenation, resulting in blocks lasting approximately 3 minutes each. All stories were normalised to the same root mean square sound amplitude. To ensure acoustic comparability between attended stories in English and French, we computed the residual power spectra of the English and French speech envelopes. Analyses revealed no significant differences between the two languages: both sets of envelopes exhibited residual power between 1 and 9 Hz, with a prominent peak around 4–5 Hz (Figure 1B); consistent with the temporal modulations typically reported in natural speech (Chandrasekaran et al., 2009; Peelle et al., 2013).

### 2.4 Procedure

Participants first completed a short practice trial to familiarise themselves with the task. They were then presented with 16 blocks, each involving listening to the attended story while ignoring the unattended stream, followed by comprehension questions. The attended ear alternated between blocks. Each block began with a recorded auditory cue indicating which ear participants should attend to. Block order was pseudo-randomised in two ways. We first counterbalanced which language was attended first (English or French) and this remained constant throughout the experiment to avoid language switching effects (i.e. either all English conditions were presented first, or all French conditions). Next, the type of interference (MuR or speech) was counterbalanced and matched between language conditions (e.g., participants starting with MuR in English also started with MuR in French). Finally, we counterbalanced which of the two stories within each condition was presented first. To ensure that participants were paying attention, they were asked to listen attentively to the instructed side, and told they will be completing a set of comprehension questions after each block. There were ten questions after each block, for a total of 160 responses per participant. Participants answered the comprehension questions on a laptop. No feedback was provided. The experiment took approximately 1 hour and 30 minutes.

### 2.5 Behavioural data

The comprehension task consisted of 40 multiple-choice questions per condition (160 in total), with four answer options for each question. Additionally, participants were required to indicate their confidence in each answer by stating whether they were certain, unsure, or guessing. Overall, participants answered the questions with an average accuracy of 90.8% (SD = 6.2), and their average guessing rate was 3.0% (SD = 4.7), indicating that they understood the stories and were able to perform the dichotic listening task effectively.

Five questions were removed from the analysis: four due to their difficulty (average response accuracy of 50% of lower) and one because of incorrect wording, retaining 155 answers per participant. Comprehension accuracy was modelled using Generalised Linear Mixed-Effects Models in RStudio (version 4.2.2, 2022). The data were fitted with the fixed effects of Usage Group (Active, Moderate, Inactive), Condition (English-MuR, English-English, French-MuR, and French-French), and the control variables of L2 AoA, L2 Proficiency, and number of Languages Spoken. Random intercepts for Participants and Questions were included. Contrasts were sum-to-zero contrast coded. P-values were calculated using the Type III sum of squares method with the Satterthwaite approximation for degrees of freedom, provided by the Anova function of the ‘car’ package (version 3.1-2, Fox et al., 2012). Post-hoc tests were performed using the ‘emmeans’ package (version 1.8.6, Lenth et al., 2019) applying a Tukey correction. Significant p-values are reported at the .05 threshold.

### 2.6 EEG data acquisition and pre-processing

A high-density 128 channel electrodes cap (Electrical Geodesics Inc., Eugene, OR, USA) was connected to the EGI Net-Station software (version 5.4.2) via a GES 400 amplifier. Impedance for each electrode was kept under 100Ω. 36 channels located in the peripheral areas of the net were excluded to reduce muscle artifacts. The data were preprocessed in MATLAB using the EEGLAB toolbox (Delorme & Makeig, 2004; version 2024.2). A bandpass filter from 1 to 40 Hz was applied to the continuous data, which was subsequently down-sampled to 100 Hz. Bad channels were identified using automatic detection based on kurtosis (±5 SDs) and power spectrum (±3 SDs); all flagged channels were visually inspected and manually corrected where necessary. After interpolation of the bad channels and re-referencing to the average of all channels, an Independent Component Analysis (ICA) was performed for artefact detection. ICA components were automatically classified using the ICLabel plugin (Pion-Tonachini et al., 2019), and confirmed or adjusted by visual inspection. Data were epoched 0–2400 ms from sentence onset. The first epoch of each block was excluded to account for potential attentional settling effects. Bad epochs were automatically rejected using the pop_autorej tool, applying amplitude and joint-probability thresholds (±3 SD), resulting in a final rejection rate of 8.87%, consistent with recommended standards (Delorme et al., 2007).

### 2.7 mTRF analysis

The multivariate Temporal Response Function (mTRF) analysis was conducted to measure neural tracking of the speech envelope, using the mTRF Toolbox in MATLAB (Crosse et al., 2016). Speech envelopes were extracted using the ‘Hilbert2’ function from EEGLAB and downsampled to 100 Hz to match the EEG sampling rate. EEG data were z-score normalised and bandpass filtered between 1 and 12 Hz to attenuate high-frequency noise. Epochs from the preprocessed EEG data were concatenated for each participant and condition, with 100 ms of zero-padding between epochs to prevent edge artefacts. The same procedure was applied to the corresponding speech envelopes.

Backward (decoding) mTRF analysis was used to estimate how accurately the speech envelope could be reconstructed from the EEG signal. Models were computed on approximately 10 minutes of continuous listening data using a 10-fold cross-validation procedure, with a time-lag window from 0 to 600 ms. The resulting Pearson correlation coefficients (r-values) between the predicted and actual speech envelopes were averaged across folds. Statistical significance was assessed using linear mixed-effects regression models (lmer) in R, applied to the per-participant, per-condition r-values. The p-values were considered significant at p < .05. In addition to computing the backward models across the broadband spectrum (1-12Hz), backward models in delta (1–4 Hz), theta (4–8 Hz) and alpha (8–12 Hz) frequency bands were also computed. Frequency decomposition was performed using MNE-Python, applying an 8th-order Butterworth IIR filter with zero-phase filtering before exporting the filtered data back to EEGLAB format. To estimate chance-level performance against which model performance in each frequency band could be evaluated, a permutation approach was used in which the speech envelope was circularly shifted by a randomised lag, disrupting the temporal alignment between stimulus and response while preserving the overall acoustic structure. For each mTRF model, 100 r-values were generated from permuted data, resulting in 400 control r-values per participant across the four conditions. The actual r-values were compared to the chance-level r-values using paired-sample t-tests. In each condition, for both attended and unattended speech, broadband, Delta, and Theta r-values were all significantly higher than their respective control values (Appendix, Table A-1), confirming reliable neural tracking above chance level. R-values in the Alpha band were however not consistently different from control values across conditions in either attended or unattended streams, and were therefore not further analysed. Following Crosse et al. (2021), we also calculated d-prime scores by comparing the actual r-values to those from the permuted models for each participant. The average d-prime was 2.69 (SD = 1.02), indicating robust model performance well above the commonly used reliability threshold. A Kruskal–Wallis test showed no significant difference in d-prime between Usage groups (χ²(2) = 0.26, p = .879).

### 2.8 Power Spectral Density analysis

Power Spectral Density (PSD) analysis was performed to measure oscillatory power in specific frequency bands during listening, using the FieldTrip toolbox (Oostenveld et al., 2011) in MATLAB. The baseline (extracted from 10 sec pre-listening period in each block) and listening periods were each divided into 2.4-second epochs to ensure equivalent segment lengths and PSD was computed for each participant and condition, matching the same segmentation scheme used in the mTRF analysis. PSD was computed using the multitaper frequency transformation method (mtmfft) with a Hanning taper, calculated across the full epoch length in 1Hz steps from 1 to 12 Hz. Power spectra were averaged across channels and time, and a baseline-corrected (normalised) power spectrum was obtained by subtracting the baseline power from the corresponding listening period spectrum. Band power was computed as the mean power within each frequency band (delta: 1-4 Hz, theta: 4-8 Hz, alpha: 8-12 Hz). Statistical analyses were conducted using linear mixed-effects models (lmer), mirroring mTRF analyses.

## 3. Results

### 3.1 Behavioural Data

To examine whether bilinguals’ language usage modulates their accuracy on comprehension questions across conditions, the first model included the fixed effects of Usage Group (Active, Moderate, Inactive), Condition (English-MuR, English-English, French-MuR, and French-French), and their interaction, and relevant control variables (L2 AoA, L2 Proficiency, Multilingualism). Results revealed significant effects of Condition (χ²(3) = 22.67, p < .001), and a significant Usage Group × Condition interaction (χ²(6) = 33.06, p < .001), suggesting differential effects of Usage across Conditions (Figure 1C and Table A-2a). L2 AoA also emerged as a significant predictor (χ²(1) = 7.80, p = .005). To unpack the Usage Group × Condition interaction and asses if it was driven by the attended language (L1 or L2) or interference type (intelligible or unintelligible), we ran a model with the predictors Usage Group, Attended Language (English vs French), Interference Type (Musical Rain vs Language), and their interactions. There was a significant effect of Attended Language (χ²(1) = 21.16, p < .001), with better comprehension in English than in French conditions; as well as a significant Usage Group × Attended Language interaction (χ²(2) = 8.06, p = .018), revealing that the English language advantage was smallest in the Active-Usage group (Figure 1D). There was also a significant interaction between Usage Group and Interference Type (χ²(2) = 22.72, p < .001), with the Inactive-Usage group performing significantly worse under intelligible (linguistic) interference (z = 3.12, *p* = .002), and Moderate- and Active-Usage groups not affected by the Interference Type (Figure 1D and Table A-2b). Since L2 AoA also emerged as a significant predictor, we confirmed that all these effects remained robust when taking AoA into account (Table A-2c). Overall, the behavioural findings suggest that usage modulates comprehension performance in a dichotic listening task, with inactive bilinguals showing least balanced comprehension across their L1 and L2, as well as weakest resistance to intelligible interference.

#### 3.2.1 mTRF analysis: Broadband (1-12 Hz)

Linear mixed effects were used to model r-values between decoded and actual speech envelopes and examine how bilinguals from different usage groups tracked attended and unattended streams across conditions. The first and most general model included the fixed effects of Attention (Attended vs Unattended), Condition, Usage Group, and their interactions. This model revealed a robust effect of Attention (F(1, 315) = 450.05, p < .001, η^2^ = .59), with higher r-values (i.e. better decoding) in the attended compared to the unattended stream. There was also an effect of Condition (F(3, 315) = 11.53, p < .001, η^2^ = .10), and a significant Attention × Condition interaction (F(3, 315) = 24.91, p < .001, η^2^ = .19), indicating that attentional effects varied by condition. Usage Group was not a significant predictor (F(2, 45) = 0.02, p = .980). See Figure 2A and Table A-3a for full details of this analysis. Given the significant effect of AoA on behavioural data, we also replicated these analyses to include AoA as a predictor. Results replicated the findings reported above, and showed no significant effects of AoA on speech decoding (Table A-3b).

**Figure 2.**
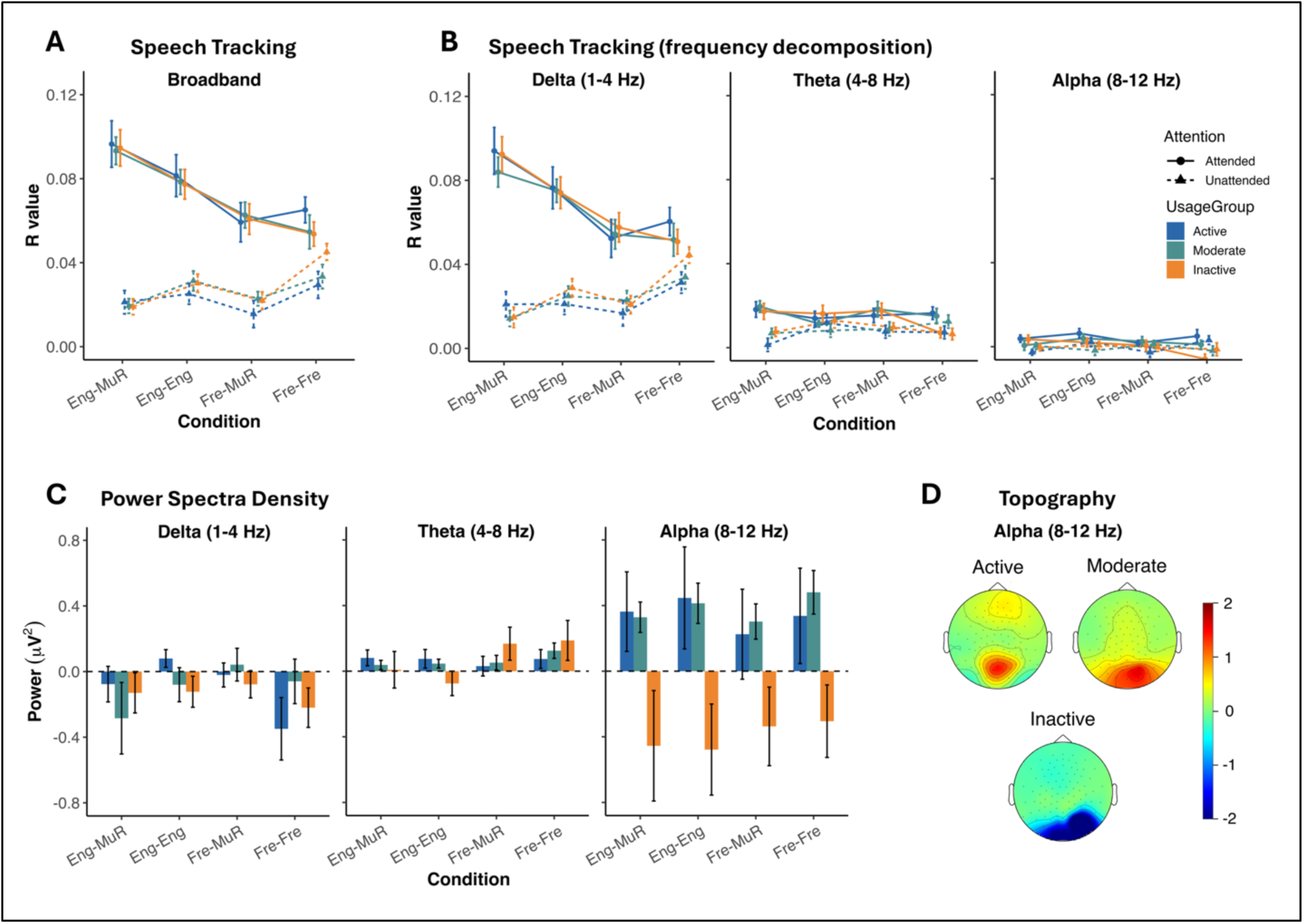
EEG results for speech tracking and power spectrum density across frequency bands. **(A) Speech tracking (broadband 1–12 Hz)**, mTRF decoding for attended and unattended speech across groups, showing effects of attention, condition, but no effect of group. **(B) Speech tracking by frequency band (Delta 1–4 Hz, Theta 4–8 Hz, Alpha 8-12 Hz)**, showing that most tracking occurred in the Delta band, with weaker effects in Theta and no reliable tracking in Alpha. **(C) Power Spectral Density (PSD)**, baseline-corrected power during listening across Delta, Theta, and Alpha bands, showing an overall reduction in Delta power, increased Theta power when attending to French (L2) particularly in the Inactive group, and increased Alpha power in Active and Moderate users but decreased Alpha power in the Inactive group. **(D) Alpha topography**, scalp distribution of Alpha power, showing that the observed increases and decreases originate from parietal regions. Error bars represent ± SEM.

To better understand the effects of Condition on the decoding of Attended and Unattended streams, we applied the same approach as in the behavioural analyses above and analysed them as a function of Attended Language and Interference Type, as well as the Usage Group. This analysis was run separately for Attended and Unattended streams. For Attended stream (see Table A-4), we found a significant effect of Attended Language (F(1, 135) = 78.99, p < .001, η^2^ = .37), with stronger decoding of English compared to French – mirroring the results of the behavioural analyses. Interference Type also significantly affected decoding of the attended stream (F(1, 135) = 9.21, p = .003, η^2^ = .06), with weaker decoding of attended streams paired with intelligible interference. There was also a significant interaction between Attended Language and Interference Type (F(1, 135) = 4.15, p = .044, η^2^ = .03), such that the effect of intelligibility was only significant when listening to English (z = 3.59, p < .001). As in the general model above, we found no effect of Usage Group (F(2, 45) = 0.11, p = .892). For the Unattended stream (see Table A-4), decoding was significantly influenced by Interference Type (F(1, 135) = 26.56, p < .001, η^2^ = .16), with stronger tracking of intelligible interference (language) compared to unintelligible interference (Musical Rain). There were no effects of Attended Language (F(1, 135) = 2.28, p = .133), nor any interactions between them (F(1, 135) = 2.01, p = .159). Usage Group was again not a significant predictor (F(2, 45) = 0,97, p = .388).

Together, these results suggest stronger decoding of the attended stream compared to the unattended stream, replicating a well-established result from the literature (Ding & Simon, 2012; O’Sullivan et al., 2015; Rimmele et al., 2015). Tracking of the attended speech was modulated by both the attended language (with L1 English decoded better than L2 French) and the type of the interference (with stronger decoding of attended stream paired with unintelligible interference, particularly in participants’ L1); while tracking of the unattended stream was only modulated by the intelligibility of that stream. None of these processes was affected by the degree of L2 usage in bilinguals.

#### 3.2.2 mTRF analysis: Frequency decomposition

To characterise the nature of the observed effects and their likely source, we examined frequency-specific decoding in the frequency bands that were consistently above chance level - Delta (1-4 Hz) and Theta (4-8 Hz). Following the same procedure as described above, in each frequency band we first ran a general model (including the fixed effects of Attention, Condition, Usage Group, and their interactions), followed by models testing for effects of Attended Language and Interference Type. General models in both Delta and Theta frequency bands showed a significant effect of Attention, with higher decoding of the attended than the unattended stream (Delta: F(1, 315) = 384.15, p < .001, η^2^ = .55; Theta: F(3, 315) = 33.37, p < .001, η^2^ = .10), as well as a significant Attention × Condition interaction (Delta: F(1, 315) = 27.80, p < .001, η^2^ = .21; Theta: F(3, 315) = 3.52, p = .016, η^2^ = .03).

Significant main effect of Condition only emerged in the Delta band however (Delta: F(3, 315) = 9.46, p < .001, η^2^ = .08; Theta: F(3, 315) = 0.55, p = .651). Usage Group did not emerge as a significant predictor in either frequency band (Delta: F(2, 45) = 0.12, p = .885; Theta: F(2, 45) = 0.16, p = .855). See Figure 2B and Table A-5 for the full results.

Analysis of the attended stream in the Delta band (Figure 2B, Table A-6) replicated all equivalent broadband effects. We found a significant effect of Attended Language (F(1, 135) = 80.40, p < .001, η^2^ = .37), with stronger tracking of English than French. There was also a significant effect of Interference Type (F(1, 135) = 6.01, p = .016, η^2^ = .04), with reduced tracking of the attended stream when it was paired with intelligible interference. Finally, we observed a significant Attended Language × Interference Type interaction (F(1, 135) = 5.35, p = .022, η^2^ = .04), with the interference effect only emerging in the English conditions (z = 3.39, p < .001). These findings suggest that Delta band effects drive the overall speech tracking results found across the broadband spectrum, likely reflecting the effortful listening conditions inherent in the dichotic listening task. In the analysis of the attended stream in the Theta band (Figure 2B, Table A-6), we only found a significant effect of Interference Type (F(1, 135) = 6.58, p = .011, η^2^ = .05), with weaker decoding of the attended stream paired with intelligible interference. However, there was no significant effect of Attended Language, nor any interaction between the two. The finding that only Interference Type affected cortical tracking in theta band is in line with previous literature (Doelling et al., 2014; Mai et al., 2016; Peelle et al., 2013) that suggests that theta band oscillations predominantly track low-level speech features through bottom-up mechanisms, which are more strongly disrupted by intelligible (linguistic) interference. Analyses of the unattended streams showed comparable results to those seen for the attended signal, with the Delta band results largely replicating broadband analyses, and Theta band results not revealing any significant effects – see Table A-6 for details. The results of these frequency-specific analyses suggest that they can indeed differentiate between the bottom-up tracking of the signal in theta band and the processes associated with effortful speech processing in delta band. None of these processes were however modulated by the degree of language usage.

### 3.3 Power Spectrum Density (PSD) analysis

The final set of analyses focused on frequency-specific power changes from baseline across usage groups and conditions in the Delta (1–4 Hz), Theta (4–8 Hz), and Alpha (8–12 Hz) frequency bands, with the latter one in particular potentially indicative of task demands and resulting top-down resource allocation during our listening task.

In the Delta band, a paired-sample t-test between power during listening and power during baseline revealed an overall significant reduction in power during the task (t(191) = -2.99, p = .003). Analysis of the baseline-corrected power (power during resting state minus power during task) across groups and conditions showed no significant effects of Condition, Usage Group, or their interaction (all p<.05, Figure 2C and Table A-7). This overall suppression of Delta band power during the listening task may indicate good task engagement across groups and conditions, reflecting a transition from a resting to an active attentional state and a redistribution of neural resources toward higher-frequency oscillations (Harmony, 2013; Knyazev, 2012). In contrast, analysis of Theta power against the baseline revealed an overall increase during the task (t(191) = 3.29, p = .001). Modelling the baseline-corrected Theta power against Condition, Usage Group and their interaction revealed a significant main effect of Condition (F(3, 135) = 23.48, p = .018) and a significant Usage Group × Condition interaction (F(6, 135) = 2.57, p = .022, Figure 2C, Table A-7). With theta-power increase generally associated with increased cognitive demand (Cavanagh & Frank, 2014; Klimesch, 1999), the follow up model aimed to establish whether this increase is triggered by Usage Group interacting with Attended Language or Interference Type (see Table A-8). The results showed a significant effect of Attended Language (F(1, 135) = 8.46, p = .004), with higher theta increase when participants listened to French than to English; and a significant Attended Language × Usage Group interaction (F(2, 135) = 6.96, p = .001). Post-hoc comparisons showed that this interaction reflected higher theta power in the Inactive-Usage group when attending to French relative to English (t(135) = 4.59, p < .001), whereas no such difference was observed in the Active- or Moderate-Usage groups. Jointly these results suggest increased cognitive demand when processing an L2, particularly in Inactive-Usage bilinguals.

In the Alpha band, the paired-sample t-test showed no difference between power during listening and power during baseline (t(191) = 1.55, p = .122) but this masked a striking divergence between Usage Groups. Indeed, a model of baseline-corrected alpha power against Condition, Usage Group and their interaction (Figure 2C, Table A-7) showed a significant effect of Group (F(2, 45) = 4.23, p = .021), with the Inactive-Usage group exhibiting significantly lower alpha power than both the Moderate-Usage (z = -2.58, p = .034) and Active-Usage groups (z = -2.45, p = .047), who did not differ from each other. These findings highlight group-specific modulation of alpha activity, with alpha power decreasing in Inactive-Usage bilinguals during dichotic listening, and increasing in Moderate- and Active-Usage bilinguals, arguably reflecting a more efficient top-down control in these usage groups. Topographical plots of alpha power (Figure 2D) show that these effects were localised over parietal regions, consistent with the role of posterior alpha oscillations in supporting attentional suppression (Obleser & Weisz, 2012; Wöstmann et al., 2017; see Foxe & Snyder, 2011 for review). AoA was not a significant predictor in any of the PDS analyses (see Table A-9).

## 4. Discussion

This study investigated whether selective attention adaptations in bilinguals are dynamically modulated by current L2 usage, and what neural mechanisms might underpin this effect. We tested highly proficient English-French bilinguals who were matched for L2 proficiency and learning history but differed in their frequency of L2 usage. We examined how usage modulates comprehension, cortical speech tracking, and spectral power across frequencies while participants listened to continuous speech paired with intelligible or unintelligible interference. Behaviourally, Inactive bilinguals were most affected by the presence of intelligible (linguistic) interference, whereas Moderate- and Active-Usage groups maintained robust performance across all interference types. Cortical tracking results were mainly driven by the delta band and not modulated by L2 usage. Spectral power analyses revealed prominent usage effects in the alpha band, with Inactive bilinguals showing reduced power during the listening task relative to the other two groups. Together, these findings demonstrate that L2 usage does not affect how bilinguals encode attended speech, but instead specifically modulates the top-down inhibitory mechanisms reflected in the degree of alpha band activity. We discuss these results in more detail below.

### 4.1. Usage e[ects on selective attention in bilinguals

The primary question in this study was whether the degree of L2 usage modulates selective attention processing in bilinguals. This effect is predicted by the view that repeated selecting of the target language and suppression of the non-target one underpin effective resisting to interference, as postulated by the Adaptive Control Hypothesis (Green & Abutalebi, 2013). It is also in line with the existing evidence that modification of bilingual processing mechanisms declines with reduced language usage (Hosoda et al., 2013; Tu et al., 2015), although these changes do not necessarily follow a linear trajectory (Korenar et al., 2023).

Our data provide evidence for both usage effects on bilingual selective attention, and the non-linear nature of these changes. Behaviourally, while all participants struggled more with processing attended speech in their L2, and when this was paired intelligible interference, this was particularly prominent in Inactive-Usage bilinguals. More specifically, while Active-Usage and Moderate-Usage participants performed similarly across all conditions, a significant usage effect emerged when the influence of attended language and interference type combined to create a particularly demanding listening condition, robustly affecting Inactive-Usage bilinguals. This suggests that highly and moderately active bilinguals may be better able to resist more disruptive speech processing contexts. These findings replicate previous studies on active bilinguals (Filippi et al., 2012, 2015; Olguín et al., 2019) and underscore that even moderate usage of the second language might be sufficient to underpin this effect.

Further evidence for usage effects emerged from spectral power analyses, where Inactive bilinguals also showed a distinct pattern of results compared to the other two groups, particularly in the alpha frequency band. Here, Active-Usage and Moderate-Usage bilinguals exhibited increased alpha power during listening relative to a baseline, while Inactive-Usage bilinguals showed decreased alpha power during listening relative to a baseline. The modulation in alpha band power has been associated with inhibition and attentional control (e.g., Deng et al., 2020; Klimesch, 2012; Strauß et al., 2014), and we discuss the functional significance of this finding in the next section. Furthermore, these group differences in the alpha band power were independent of condition, suggesting a general attentional control effect rather than one tied to specific language or interference type. Here again we saw no evidence for a linear trajectory of these usage effects, with Active-Usage and Moderate-Usage bilinguals clearly patterning together, and providing further evidence that even moderate usage of L2 modulates attentional and inhibitory processes in bilinguals. Effects of usage also emerged in the theta band, which has been linked to increased cognitive demand (Cavanagh & Frank, 2014; Klimesch, 1999), particularly during language comprehension (Bastiaansen & Hagoort, 2003; Meyer, 2018). Here, Inactive-Usage bilinguals showed higher theta power when listening to narratives in their L2 (French) compared to their L1 (English), consistent with the idea that limited exposure requires additional cognitive resources to process the less frequently used language.

By contrast, results from the delta band showed power decrease across all groups relative to the resting-state baseline, suggesting uniform task engagement across participants, and likely reflecting a downregulation of default-mode processing to prioritise attentional mechanisms during dichotic-listening (Harmony, 2013; Kaushik et al., 2022; Knyazev, 2012). The fact that this reduction occurred consistently across usage groups implies that basic engagement mechanisms are (unsuprisingly) preserved regardless of recent L2 experience. No effects of usage emerged in any of the cortical tracking analyses.

Jointly, these findings highlight that the degree of L2 usage plays a significant role in modulating the mechanisms of selective attention in bilinguals, and also potentially affect their behavioural outcomes further down the line. The finding that usage affects inhibitory power activity and indices of higher cognitive load in alpha and theta bands, rather than tracking of the attended and unattended speech envelopes, is indicative of the mechanism that underpins these effects, which we discuss in the next section.

### 4.2. The neural mechanisms of L2 usage e[ects

The second question driving the current study was about the neural mechanisms that might underpin the effects of L2 usage on selective attention in bilinguals. The available literature shows robust evidence that bilingualism affects cortical tracking of attended speech, such that this is less affected by changes in interference compared to monolinguals (Olguin et al, 2019; Phelps et al, 2022). Cortical speech tracking in naturalistic listening is thought to capture both bottom-up entrainment of the neural signal to acoustic changes of the speech signal, as well as top-down modulation and predictive processing, which is particularly prominent in adverse listening conditions (Beier et al, 2021; Luo & Ding, 2020). These different but complementary processes have also been linked to different frequency bands, with cortical tracking of speech in the delta band associated with adverse listening conditions and increased reliance on higher-order syntactic and semantic processes (Klimovich-Gray et al., 2021), and theta band more commonly linked to basic auditory processing of the acoustic regularities of speech (Di Liberto et al., 2015; Kösem et al., 2016).

Multivariate Temporal Response Function (mTRF) analysis revealed robust neural tracking of speech across all bilingual groups, with stronger tracking of the attended stream compared to the unattended stream across all conditions. This is consistent with established findings on attentional enhancement in dichotic listening tasks (Ding & Simon, 2012; O’Sullivan et al., 2015; Rimmele et al., 2015), clearly indicating that cortical tracking is sensitive to attentional manipulations and is a robust indicator of the underlying attentional processes. Attended language (L1 or L2) and type of interference (intelligible or unintelligible) also robustly affected cortical tracking of speech, closely replicating the behavioural results described above. The finding that the decoding of narratives in participants’ L1 was consistently stronger than in their L2 – irrespective of the degree of L2 usage or age of acquisition – is in line with findings that native-language speech sounds, phonemic distinctions, and prosodic and acoustic cues tend to be encoded more precisely, robustly, or with greater fidelity (Chung & Bidelman, 2016; Lizarazu et al, 2021) and replicates existing evidence for stronger tracking of L1 over L2 in dichotic listening conditions (Grant et al., 2022). The type of interference effect was prominent across both attended and unattended streams, with intelligible interference increasing the tracking of the unattended stream while simultaneously reducing the tracking of the attended stream. Despite this effect being attenuated in bilinguals compared to monolinguals (Olguin et al, 2018, 2019; Phelps et al, 2022), such distribution is fully expected, suggesting that gain in distractor’s intelligibility degrades the representation of attended speech. This is likely due to the linguistic competition between the two intelligible streams, and was previously shown to be specifically prominent in the delta band (Dai et al, 2022; Har-shai & Zion Golumbic, 2021). Indeed, analyses of specific frequency bands showed clear dissociation between speech tracking in delta and theta bands, with delta band results essentially replicating those seen in the broadband analysis, and theta band results only revealing weak modulation by interference type (and no effects of the attended language). As discussed, such results in the delta band are fully consistent with the widely accepted view that they reflect higher-order processing of the sentential structure (Ding et al., 2016; Giraud & Poeppel, 2012; Molinaro & Lizarazu, 2018), particularly in adverse listening conditions (Ershaid et al., 2024; Mai & Wang, 2019). Theta band results on the other hand suggest equivalent encoding of the bottom-up, low-level acoustic features across both languages, which was expected given that speech envelopes of L1 and L2 stories in our experiment were carefully matched, with residual power distributed equally across frequencies in the two languages (Fig 1B). Together, delta and theta band effects suggest that the type of distractor affects both bottom-up processing and top-down tracking of the signal (Doelling et al., 2014; Peelle et al., 2013), as well as showing that they clearly capture two different processing aspects.

Critically, however, none of these effects were modulated by language usage. This could be potentially explained by the fact that the current study focused on the tracking of the acoustic envelope of speech, which is important for speech intelligibility (Peelle et al., 2013), but largely limited to perceptual features of speech such as rhythm, prosody, stress, and broad phonetic boundaries. Our results imply that the cortical tracking of speech at this level of encoding is similar across bilinguals, regardless of their recent L2 usage. In other words, the primarily perceptual aspects of speech processing, indexed by the tracking of the speech envelope, are likely to be stable and not shaped by recent language experience.

However, as outlined above, spectral power analysis revealed distinct neural signatures of attentional control across different usage groups during the dichotic listening task. Alpha band power in particular revealed pronounced differences between Active- and Moderate-Usage bilinguals on the one hand, and Inactive-Usage group on the other; with the former exhibiting significantly increased alpha power during listening. This is arguably indicative of Event-Related Synchronisation (ERS, Pfurtscheller & Lopes Da Silva, 1999) - a transient increase in oscillatory alpha power relative to baseline that is typically interpreted as reflecting inhibition and attentional control (Benedek et al., 2014; Deng et al., 2020; Klimesch, 2012); consistent with studies linking alpha synchronisation with top-down inhibitory control over irrelevant auditory input (Strauß et al., 2014; Wöstmann et al., 2017). Conversely, Inactive-Usage bilinguals showed decreased alpha power during listening, reflecting Event-Related Desynchronisation (ERD, Pfurtscheller & Lopes Da Silva, 1999), a neural signature typically linked to cognitive overload or weakened inhibitory control under demanding conditions (Krause et al., 2000; Obleser et al., 2012; Wisniewski et al., 2017). These findings highlight the central role of current L2 usage in sustaining the neural efficiency of top-down control processes during auditory attention tasks.

Taken together, our findings suggest that selective attention processes in bilinguals are characterised by multiple complementary mechanisms. The first mechanism pertains to lower-level perceptual tracking of speech, as captured by mTRF decoding. This process was uniformly sensitive to interference type and attended language, yet unaffected by usage group, indicating that lower-level decoding of speech operates independently of recent language experience. The second mechanism, reflected in alpha-band power, appears to index the allocation of top-down attentional control, and is strongly modulated by L2 usage. This combination of processes arguably provides a mechanistic explanation for the selective behavioural vulnerability of Inactive-Usage bilinguals. While interference disrupted neural tracking in all participants, only those with reduced alpha power suffered a corresponding decline in comprehension. In contrast, Active- and Moderate-Usage bilinguals showed increased alpha synchronisation, consistent with efficient inhibitory control, which may have led to preserved behavioural performance. These contrasting alpha power responses suggest a functional dissociation in attentional regulation across usage groups: for those actively engaging with their L2, alpha ERS may reflect more effective suppression of irrelevant input, in line with predictions from the Adaptive Control Hypothesis (Green & Abutalebi, 2013). Conversely, the ERD pattern observed in Inactive-Usage bilinguals may indicate weakened inhibitory control under competing demands, potentially due to a deterioration of previously acquired attentional adaptations or due to a lack of maintained L2 suppression in daily life. Taken together, our results highlight that behavioural resilience in the face of interference is not necessarily dependent on lower-level percpetual encoding of speech, but rather on the capacity to recruit higher-level attentional control mechanisms; processes that are dynamically maintained by bilinguals’ L2 usage.

### 4.3 Dynamic Functional Adaptation in Bilinguals

In addition to showing that usage modulates selective attention adaptation in bilinguals, and revealing a possible mechanism for this, our results revealed further two key aspect that add to the literature on bilingual adaptation in selective attention. First, they suggest that selective attention adaptations are language-independent and likely supported by domain-general mechanisms of control. Although decoding of the L1 was overall higher than the L2, the key results (the modulation of comprehension by interference and usage group, the disruption of mTRF tracking by interference, and the alpha-band attentional responses modulated by usage group) were comparable when participants attended to their L1 and L2. This consistency suggests that the underlying adaptive mechanisms generalise across languages and are not tied to language-specific processes, but rather reflect broader attentional control mechanisms shaped by bilingual experience.

Moreover, the modulation by L2 usage appears to follow a non-linear pattern. Moderate-Usage bilinguals, who engaged with their L2 only several times per week, displayed alpha power and comprehension performance comparable to Active-Usage bilinguals, who use their L2 daily. This indicates that even modest but consistent L2 use may suffice to maintain the benefits of bilingual attention adaptations. In contrast, complete disengagement, characteristic of Inactive-Usage bilinguals was associated with differences in both neural and behavioural measures. This threshold-like profile echoes structural findings showing that bilingual brain adaptations may emerge or deteriorate rapidly depending on experience (Hosoda et al., 2013; Tu et al., 2015). Our results suggest that selective attention mechanisms are similarly dynamic: robust to some degree of reduction in language usage, but sensitive to complete disuse. This view is further supported by recent work showing plateau effects in bilingual cognitive adaptations (Korenar et al., 2023), and aligns with the broader understanding of neuroplasticity as flexibly shaped by experience.

These findings have important implications for the ongoing debates on the neurocognitive consequences of bilingualism. Rather than assuming a linear dose–response relationship between language use and neurocognitive outcome, they suggest that some language-driven adaptations may stabilise once a sufficient level of language engagement is reached, but degrade if usage falls below a critical minimum. Such a flexible framework of multi-faceted functional adaptation in bilinguals not only better captures the lived experience of bilinguals, but may also be instrumental in reconciling existing inconsistencies in the bilingualism literature where usage has not been taken into account.

### 4.4 Conclusions

This study sought to answer whether active language usage shapes the adaptation of selective attention in bilingualism; and what neural mechanisms might drive this process. While we saw no effects of usage on lower-level perceptual encoding of speech, oscillatory activity in alpha band was strongly modulated by usage, with active L2 use leading to more efficient inhibitory control and stronger behavioural resilience to interference. These effects were language-independent and non-linear, with even modest degree of L2 usage sufficient to maintain the efficient inhibitory control. This underlines the dynamic nature of selective attention adaptation in bilingualism, with language usage a paramount in shaping this adaptation in different contexts and throughout the life span.

## Data availability

The datasets and code for this study are available on OSF: https://osf.io/t4e2y/

## Acknowledgements

This research was supported by funds from the Department of Psychology, University of Cambridge to S.T-G. We would like to thank the participants for making this research possible.

## APPENDIX

**Table A-1:** Comparison of mTRF r values against chance *( *** p<.001; ** p<.01; * p<.05)*

| Frequency | Attention | Condition |  |  |  |
| --- | --- | --- | --- | --- | --- |
|  |  | EngMuR | EngEng | FreMuR | FreFre |
| Broadband | Attended | t(47) = 19.03 *** | t(47) = 18.04 *** | t(47) = 13.48 *** | t(47) = 15.3 *** |
|  | Unattended | t(47) = 7.64 *** | t(47) = 10.91 *** | t(47) = 6.70 *** | t(47) = 10.61 *** |
| Delta | Attended | t(47) = 17.30 *** | t(47) = 16.74 *** | t(47) = 12.61 *** | t(47) = 13.96 *** |
|  | Unattended | t(47) = 5.38 *** | t(47) = 9.43 *** | t(47) = 7.01 *** | t(47) = 12.71 *** |
| Theta | Attended | t(47) = 9.21 *** | t(47) = 7.69 *** | t(47) = 8.20 *** | t(47) = 6.51 *** |
|  | Unattended | t(47) = 3.22 ** | t(47) = 5.59 *** | t(47) = 4.97 *** | t(47) = 4.44 *** |
| Alpha | Attended | t(47) = 2.33 * | t(47) = 2.88 ** | t(47) = 1.15 | t(47) = 0.11 |
|  | Unattended | t(47) = -1.35 | t(47) = 0.36 | t(47) = -0.21 | t(47) = -0.46 |

**Table A-2:**
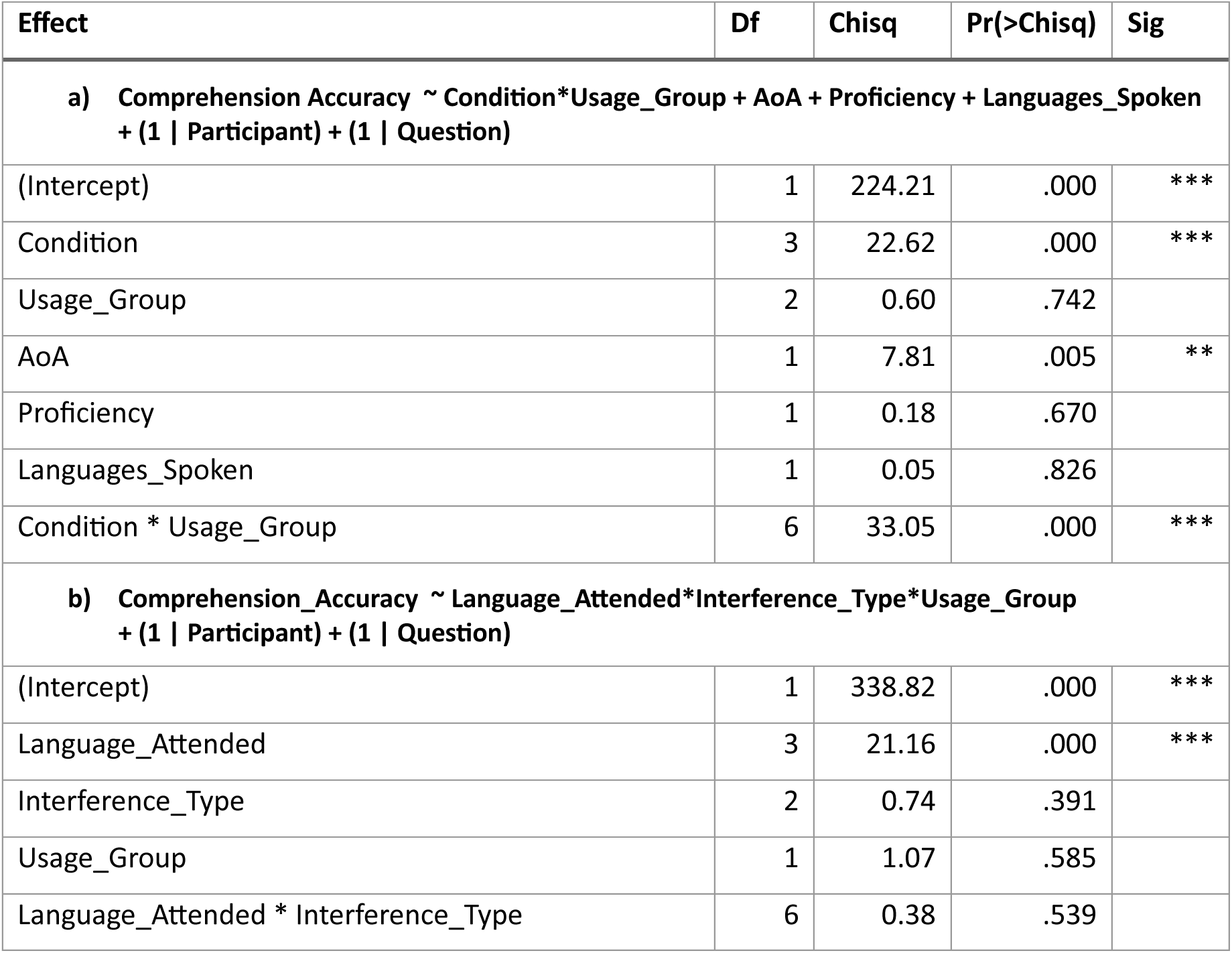
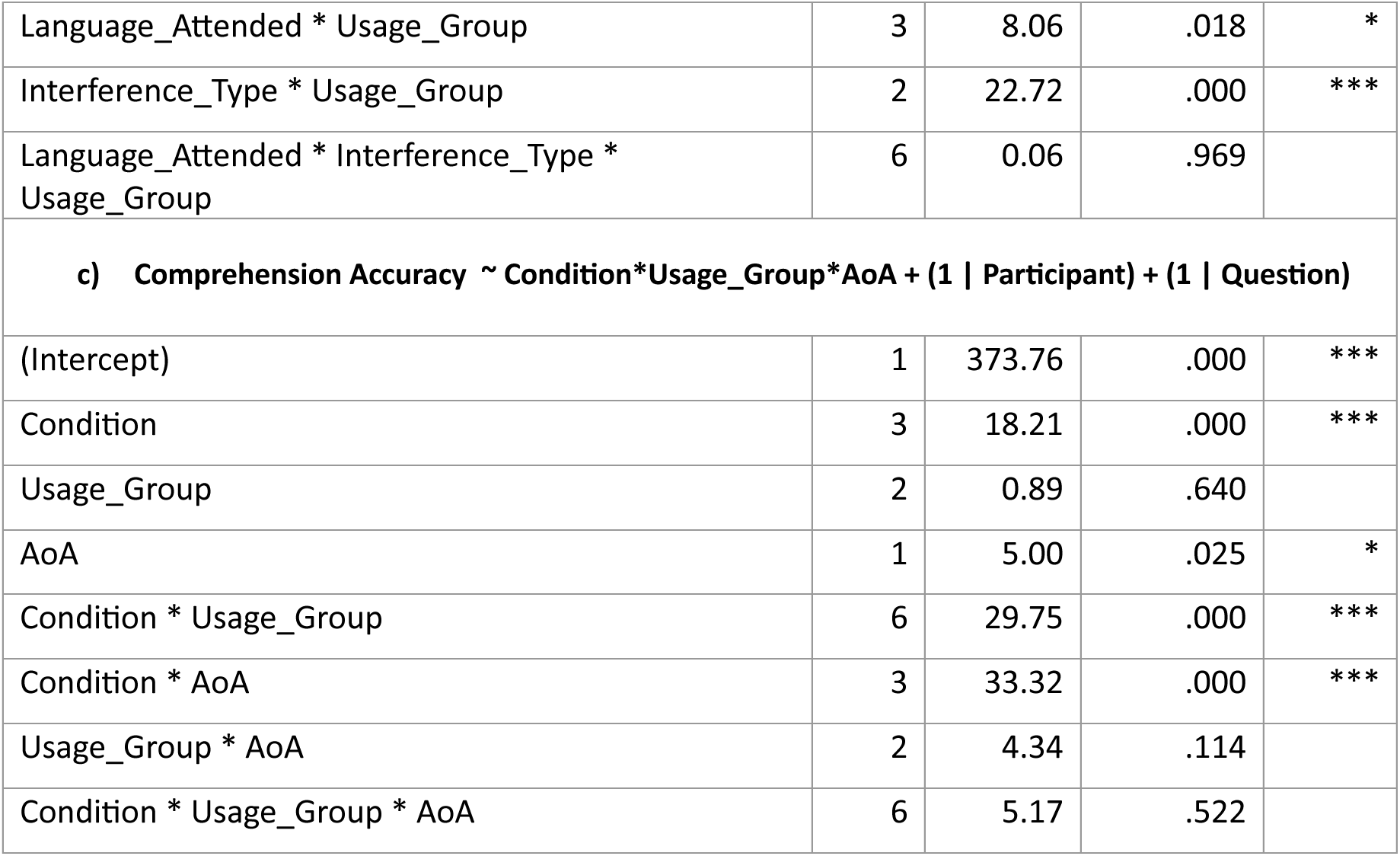
Behavioural analyses

**Table A-3:**
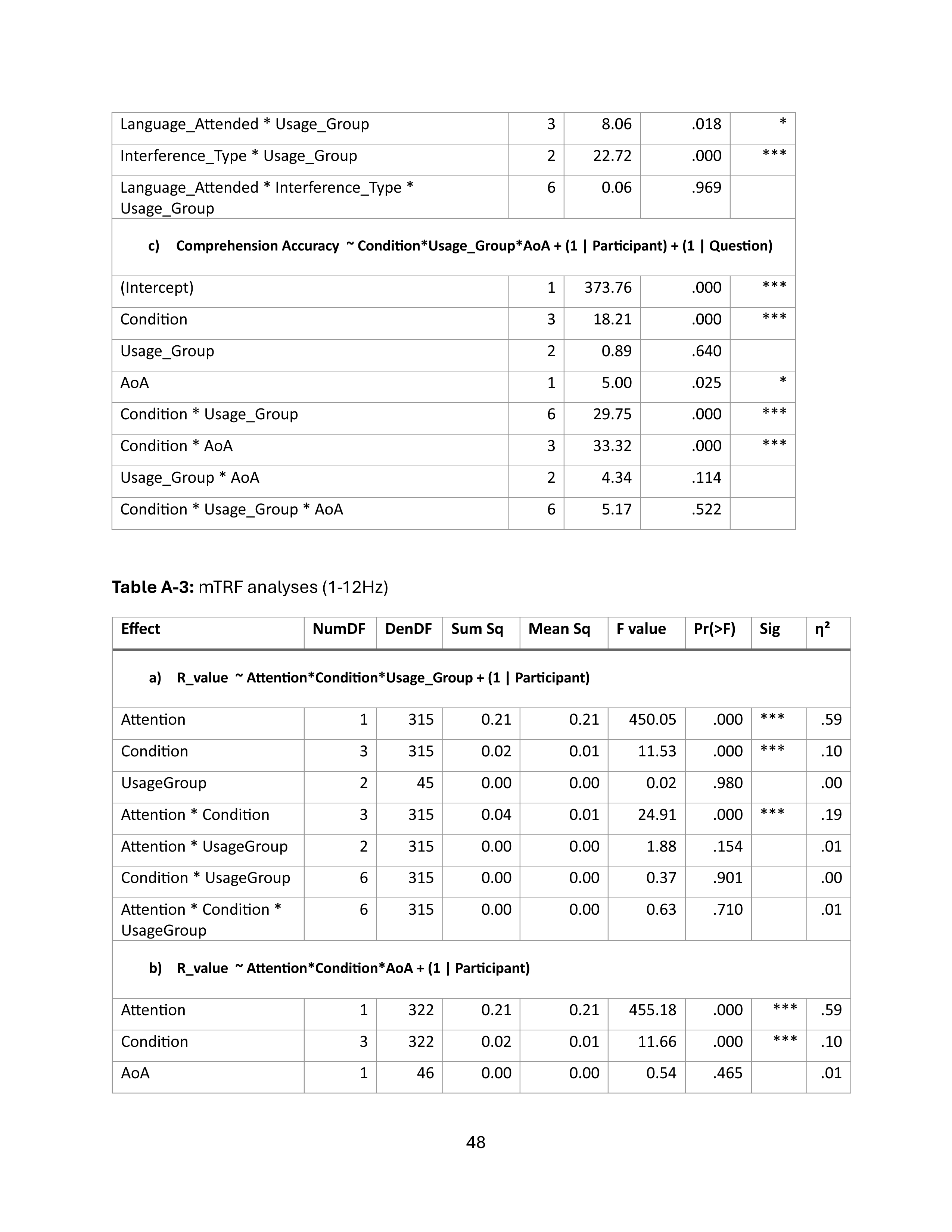
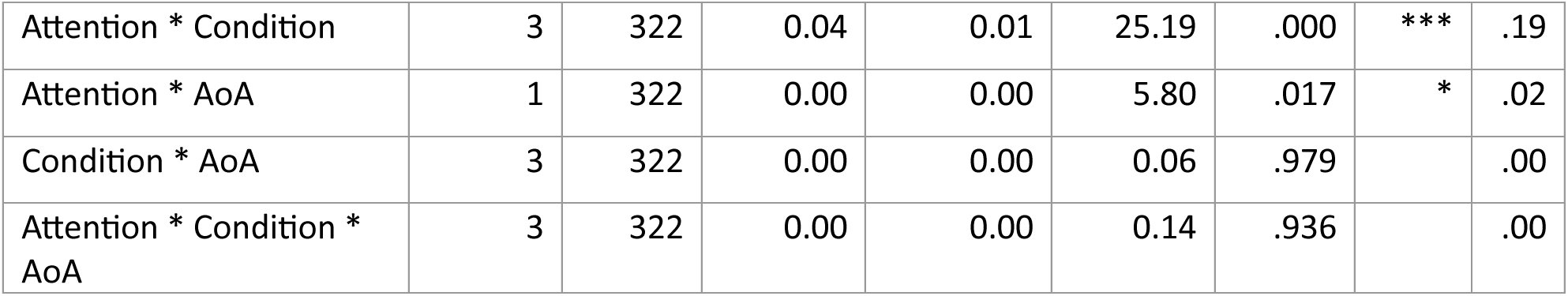
mTRF analyses (1-12Hz)

**Table A-4:** mTRF analyses (1-12Hz) for Attended and Unattended streams separately *R_value ∼ Language_Attended*Interference_Type*Usage_Group + (1 | Participant)*

| Effect | Num<br>DF | Den<br>DF | Sum<br>Sq | Mean<br>Sq | F<br>value | Pr(>F) | Sig | η² | Sum<br>Sq | Mean<br>Sq | F<br>value | Pr(>F) | Sig | η² |
| --- | --- | --- | --- | --- | --- | --- | --- | --- | --- | --- | --- | --- | --- | --- |
|  |  |  | Attended |  |  |  |  |  | Unattended |  |  |  |  |  |
| Language_Attended | 1 | 135 | 0.04 | 0.04 | 78.99 | .000 | *** | .37 | 0.00 | 0.00 | 2.28 | .133 | *** | .02 |
| Interference_Type | 1 | 135 | 0.00 | 0.00 | 9.21 | .003 | ** | .06 | 0.01 | 0.01 | 26.56 | .000 | ** | .16 |
| Usage_Group | 2 | 45 | 0.00 | 0.00 | 0.11 | .892 |  | .00 | 0.00 | 0.00 | 0.97 | .388 |  | .04 |
| Language_Attended<br>* Interference_Type | 1 | 135 | 0.00 | 0.00 | 4.15 | .044 | * | .03 | 0.00 | 0.00 | 2.01 | .159 | * | .01 |
| Language_Attended<br>* Usage_Group | 2 | 135 | 0.00 | 0.00 | 0.04 | .961 |  | .00 | 0.00 | 0.00 | 1.34 | .265 |  | .02 |
| Interference_Type *<br>Usage_Group | 2 | 135 | 0.00 | 0.00 | 0.61 | .546 |  | .00 | 0.00 | 0.00 | 1.05 | .354 |  | .02 |
| Language_Attended<br>* Interference_Type<br>* Usage_Group | 2 | 135 | 0.00 | 0.00 | 0.47 | .625 |  | .00 | 0.00 | 0.00 | 0.74 | .477 |  | .01 |

**Table A-5:** mTRF analyses (Delta and Theta frequency bands) *R_value ∼ Attention*Condition*Usage_Group + (1 | Participant)*

| Effect | NumDF | DenDF | Sum Sq | Mean Sq | F value | Pr(>F) | Sig | $\eta^2$ |
| --- | --- | --- | --- | --- | --- | --- | --- | --- |
| **<u>Delta frequency band</u>** |  |  |  |  |  |  |  |  |
| Attention | 1 | 315 | 0.19 | 0.19 | 384.15 | .000 | *** | .55 |
| Condition | 3 | 315 | 0.01 | 0.01 | 9.46 | .000 | *** | .08 |
| UsageGroup | 2 | 45 | 0.00 | 0.00 | 0.12 | .884 |  | .00 |
| Attention * Condition | 3 | 315 | 0.04 | 0.01 | 27.80 | .000 | *** | .21 |
| Attention * UsageGroup | 2 | 315 | 0.00 | 0.00 | 0.90 | .408 |  | .00 |
| Condition * UsageGroup | 6 | 315 | 0.00 | 0.00 | 0.53 | .787 |  | .00 |
| Attention * Condition * UsageGroup | 6 | 315 | 0.00 | 0.00 | 0.68 | .665 |  | .01 |

| **Theta frequency band** |  |  |  |  |  |  |  |  |
| --- | --- | --- | --- | --- | --- | --- | --- | --- |
| Attention | 1 | 315 | 0.01 | 0.01 | 33.37 | .000 | *** | .10 |
| Condition | 3 | 315 | 0.00 | 0.00 | 0.55 | .651 |  | .00 |
| UsageGroup | 2 | 45 | 0.00 | 0.00 | 0.16 | .855 |  | .00 |
| Attention * Condition | 3 | 315 | 0.00 | 0.00 | 3.52 | .015 | * | .03 |
| Attention * UsageGroup | 2 | 315 | 0.00 | 0.00 | 0.70 | .000 |  | .00 |
| Condition * UsageGroup | 6 | 315 | 0.00 | 0.00 | 1.65 | .133 |  | .03 |
| Attention * Condition * UsageGroup | 6 | 315 | 0.00 | 0.00 | 0.39 | .886 |  | .00 |

**Table A-6:**
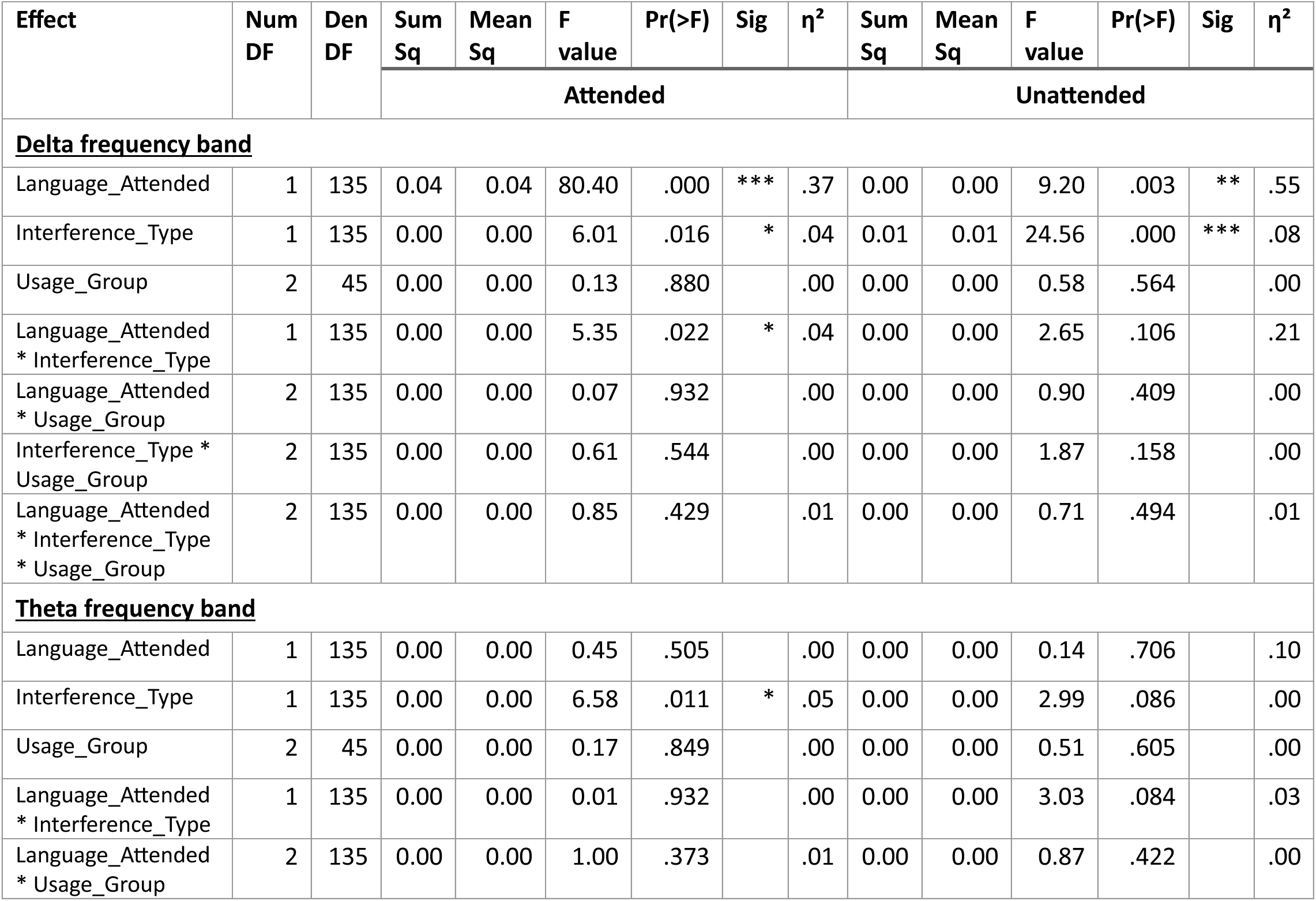
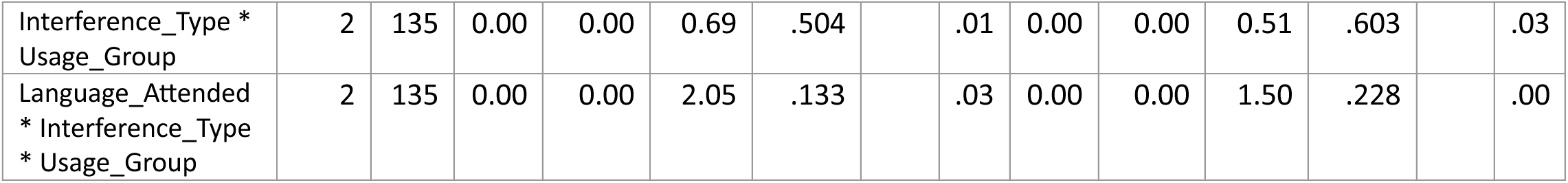
mTRF analyses (Delta and Theta frequency bands) for Attended and Unattended streams separately *R_value ∼ Language_Attended*Interference_Type*Usage_Group + (1 | Participant)*

**Table A-7:** PSD analyses (Delta, Theta and Alpha frequency bands) *Power ∼ Condition*Usage_Group + (1 | Participant)*

| Effect | NumDF | DenDF | Sum Sq | Mean Sq | F value | Pr(>F) | Sig | $\eta^2$ |
| --- | --- | --- | --- | --- | --- | --- | --- | --- |
| **<u>Delta frequency band</u>** |  |  |  |  |  |  |  |  |
| Condition | 3 | 135 | 1.24 | 0.41 | 2.61 | .054 |  | .00 |
| Usage_Group | 2 | 45 | 0.03 | 0.01 | 0.08 | .924 |  | .00 |
| Condition * Usage_Group | 6 | 135 | 1.40 | 0.24 | 1.51 | .179 |  | .00 |
| **<u>Theta frequency band</u>** |  |  |  |  |  |  |  |  |
| Condition | 3 | 135 | 0.35 | 0.12 | 3.48 | .018 | * | .07 |
| Usage_Group | 2 | 45 | 0.00 | 0.00 | 0.01 | .995 |  | .00 |
| Condition * Usage_Group | 6 | 135 | 0.50 | 0.09 | 2.57 | .022 | * | .10 |
| **<u>Alpha frequency band</u>** |  |  |  |  |  |  |  |  |
| Condition | 3 | 135 | 0.34 | 0.11 | 0.52 | .670 |  | .01 |
| Usage_Group | 2 | 45 | 1.86 | 0.93 | 4.23 | .021 | * | .16 |
| Condition * Usage_Group | 6 | 135 | 0.73 | 0.12 | 0.55 | .768 |  | .02 |

**Table A-8:** PSD analyses (Theta frequency band) *Power ∼ Language_Attended*Interference_Type*Usage_Group + (1 | Participant)*

| Effect | NumDF | DenDF | Sum Sq | Mean Sq | F value | Pr(>F) | Sig | $\eta^2$ |
| --- | --- | --- | --- | --- | --- | --- | --- | --- |
| Language_Attended | 1 | 135 | 0.29 | 0.29 | 8.46 | .004 | ** | .06 |
| Interference_Type | 1 | 135 | 0.00 | 0.00 | 0.12 | .727 |  | .00 |
| Usage_Group | 2 | 45 | 0.00 | 0.00 | 0.01 | .995 |  | .00 |
| Language_Attended * Interference_Type | 1 | 135 | 0.06 | 0.06 | 1.86 | .174 |  | .01 |
| Language_Attended * Usage_Group | 2 | 135 | 0.47 | 0.23 | 6.96 | .001 | ** | .09 |
| Interference_Type * Usage_Group | 2 | 135 | 0.04 | 0.02 | 0.65 | .523 |  | .00 |
| Language_Attended *<br>Interference_Type *<br>Usage_Group | 2 | 135 | 0.01 | 0.00 | 0.09 | .913 |  | .00 |

**Table A-9:** PSD analyses (Delta, Theta and Alpha frequency bands) *Power ∼ Condition*AoA + (1 | Participant)*

| Effect | NumDF | DenDF | Sum Sq | Mean Sq | F value | Pr(>F) | Sig | $\eta^2$ |
| --- | --- | --- | --- | --- | --- | --- | --- | --- |
| **<u>Delta frequency band</u>** |  |  |  |  |  |  |  |  |
| Condition | 3 | 138 | 0.28 | 0.09 | 0.57 | .634 |  | .01 |
| AoA | 1 | 46 | 0.31 | 0.31 | 1.91 | .174 |  | .04 |
| Condition * AoA | 3 | 138 | 0.39 | 0.13 | 0.79 | .499 |  | .02 |
| **<u>Theta frequency band</u>** |  |  |  |  |  |  |  |  |
| Condition | 3 | 138 | 0.15 | 0.05 | 1.38 | .253 |  | .03 |
| AoA | 1 | 46 | 0.05 | 0.05 | 1.27 | .266 |  | .03 |
| Condition * AoA | 3 | 138 | 0.01 | 0.00 | 0.12 | .951 |  | .00 |
| **<u>Alpha frequency band</u>** |  |  |  |  |  |  |  |  |
| Condition | 3 | 138 | 0.64 | 0.22 | 0.99 | .401 |  | .02 |
| AoA | 1 | 46 | 0.19 | 0.19 | 0.87 | .357 |  | .02 |
| Condition * AoA | 3 | 138 | 0.38 | 0.13 | 0.59 | .624 |  | .01 |

## References

1. Abutalebi, J., Canini, M., Della Rosa, P. A., Green, D. W., & Weekes, B. S. (2015). The neuroprotective effects of bilingualism upon the inferior parietal lobule: A Structural Neuroimaging Study in Aging Chinese Bilinguals. Journal of Neurolinguistics, 33, 3–13.

2. Anderson, J. A., Mak, L., Chahi, A. K., & Bialystok, E. Language and Social Background Questionnaire--Revised. Behavior Research Methods.

3. Anderson, J. A. E., Yurtsever, A., Fisher-Skau, O., Cherep, L. A., MacPhee, I., Luk, G., & Grundy, J. G. (2024). Examining the consistency in bilingualism and white matter research: A meta-analysis. Neuropsychologia, 195, 108801.

4. Bastiaansen, M., & Hagoort, P. (2003). Event-Induced Theta Responses as a Window on the Dynamics of Memory. Cortex, 39(4–5), 967–992.

5. Benedek, M., Schickel, R. J., Jauk, E., Fink, A., & Neubauer, A. C. (2014). Alpha power increases in right parietal cortex reflects focused internal attention. Neuropsychologia, 56, 393–400.

6. Berken, J. A., Chai, X., Chen, J.-K., Gracco, V. L., & Klein, D. (2016). Effects of Early and Late Bilingualism on Resting-State Functional Connectivity. The Journal of Neuroscience, 36(4), 1165–1172.

7. Birdsong, D., Gertken, L. M., & Amengual, M. (2012). Bilingual language profile: An easy-to-use instrument to assess bilingualism. COERLL, University of Texas at Austin.

8. Blanco-Elorrieta, E., Ding, N., Pylkkänen, L., & Poeppel, D. (2020). Understanding Requires Tracking: Noise and Knowledge Interact in Bilingual Comprehension. Journal of Cognitive Neuroscience, 32(10), 1975–1983.

9. Bozic, M., Tyler, L. K., Ives, D. T., Randall, B., & Marslen-Wilson, W. D. (2010). Bihemispheric foundations for human speech comprehension. Proceedings of the National Academy of Sciences, 107(40), 17439–17444.

10. Brungart, D. S., & Simpson, B. D. (2007). Effect of target-masker similarity on across-ear interference in a dichotic cocktail-party listening task. The Journal of the Acoustical Society of America, 122(3), 1724–1734.

11. Cavanagh, J. F., & Frank, M. J. (2014). Frontal theta as a mechanism for cognitive control. Trends in Cognitive Sciences, 18(8), 414–421.

12. Chandrasekaran, C., Trubanova, A., Stillittano, S., Caplier, A., & Ghazanfar, A. A. (2009). The Natural Statistics of Audiovisual Speech. PLoS Computational Biology, 5(7), e1000436.

13. Chung, W.-L., & Bidelman, G. M. (2016). Cortical encoding and neurophysiological tracking of intensity and pitch cues signaling English stress patterns in native and nonnative speakers. Brain and Language, 155–156, 49–57.

14. Chung-Fat-Yim, A., Bobb, S. C., Hoshino, N., & Marian, V. (2023). Bilingualism alters the neural correlates of sustained attention. Translational Issues in Psychological Science, 9(4), 409–421.

15. Crosse, M. J., Di Liberto, G. M., Bednar, A., & Lalor, E. C. (2016). The Multivariate Temporal Response Function (mTRF) Toolbox: A MATLAB Toolbox for Relating Neural Signals to Continuous Stimuli. Frontiers in Human Neuroscience, 10.

16. Crosse, M. J., Zuk, N. J., Di Liberto, G. M., Nidier, A. R., Molholm, S., & Lalor, E. C. (2021). Linear Modeling of Neurophysiological Responses to Speech and Other Continuous Stimuli: Methodological Considerations for Applied Research. Frontiers in Neuroscience, 15, 705621.

17. Dai, B., McQueen, J. M., Terporten, R., Hagoort, P., & Kösem, A. (2022). Distracting linguistic information impairs neural tracking of attended speech. Current Research in Neurobiology, 3, 100043.

18. Delorme, A., & Makeig, S. (2004). EEGLAB: an open source toolbox for analysis of single-trial EEG dynamics including independent component analysis. Journal of neuroscience methods, 134(1), 9–21.

19. Delorme, A., Sejnowski, T., & Makeig, S. (2007). Enhanced detection of artifacts in EEG data using higher-order statistics and independent component analysis. Neuroimage, 34(4), 1443–1449.

20. DeLuca, V., Rothman, J., Bialystok, E., & Pliatsikas, C. (2019). Redefining bilingualism as a spectrum of experiences that differentially affects brain structure and function. Proceedings of the National Academy of Sciences, 116(15), 7565–7574.

21. DeLuca, V., Rothman, J., Bialystok, E., & Pliatsikas, C. (2020). Duration and extent of bilingual experience modulate neurocognitive outcomes. NeuroImage, 204, 116222.

22. Deng, Y., Choi, I., & Shinn-Cunningham, B. (2020). Topographic specificity of alpha power during auditory spatial attention. NeuroImage, 207, 116360.

23. Di Liberto, G. M., O’Sullivan, J. A., & Lalor, E. C. (2015). Low-Frequency Cortical Entrainment to Speech Reflects Phoneme-Level Processing. Current Biology, 25(19), 2457–2465.

24. Ding, N., Melloni, L., Zhang, H., Tian, X., & Poeppel, D. (2016). Cortical tracking of hierarchical linguistic structures in connected speech. Nature Neuroscience, 19(1), 158–164.

25. Ding, N., & Simon, J. Z. (2012). Emergence of neural encoding of auditory objects while listening to competing speakers. Proceedings of the National Academy of Sciences, 109(29), 11854–11859.

26. Ding, N., & Simon, J. Z. (2013). Adaptive Temporal Encoding Leads to a Background-Insensitive Cortical Representation of Speech. The Journal of Neuroscience, 33(13), 5728–5735.

27. Doelling, K. B., Arnal, L. H., Ghitza, O., & Poeppel, D. (2014). Acoustic landmarks drive delta–theta oscillations to enable speech comprehension by facilitating perceptual parsing. NeuroImage, 85, 761–768.

28. Drgas, S., Blaszak, M., & Przekoracka-Krawczyk, A. (2021). The Combination of Neural Tracking and Alpha Power Lateralization for Auditory Attention Detection. Journal of Speech, Language, and Hearing Research, 64(9), 3603–3616.

29. Elder, C. (2018). Test review: Certifying French competency: The DELF tout public (B2). Language Testing, 35(4), 615–623.

30. Ershaid, H., Lizarazu, M., McLaughlin, D., Cooke, M., Simantiraki, O., Koutsogiannaki, M., & Lallier, M. (2024). Contributions of listening effort and intelligibility to cortical tracking of speech in adverse listening conditions. Cortex, 172, 54–71.

31. Filippi, R., Leech, R., Thomas, M. S. C., Green, D. W., & Dick, F. (2012). A bilingual advantage in controlling language interference during sentence comprehension. Bilingualism: Language and Cognition, 15(4), 858–872.

32. Filippi, R., Morris, J., Richardson, F. M., Bright, P., Thomas, M. S. C., Karmiloff-Smith, A., & Marian, V. (2015). Bilingual children show an advantage in controlling verbal interference during spoken language comprehension. Bilingualism: Language and Cognition, 18(3), 490–501.

33. Fox, J., Weisberg, S., Price, B., Adler, D., Bates, D., Baud-Bovy, G., & Bolker, B. (2012). car: Companion to Applied Regression. Version 2.0–18.

34. Foxe, J. J., & Snyder, A. C. (2011). The Role of Alpha-Band Brain Oscillations as a Sensory Suppression Mechanism during Selective Attention. Frontiers in Psychology, 2.

35. Fuglsang, S. A., Dau, T., & Hjortkjær, J. (2017). Noise-robust cortical tracking of attended speech in real-world acoustic scenes. NeuroImage, 156, 435–444.

36. Giraud, A.-L., & Poeppel, D. (2012). Cortical oscillations and speech processing: Emerging computational principles and operations. Nature Neuroscience, 15(4), 511–517.

37. Gracia-Tabuenca, Z., Barbeau, E. B., Kousaie, S., Chen, J.-K., Chai, X., & Klein, D. (2024). Enhanced efficiency in the bilingual brain through the inter-hemispheric cortico-cerebellar pathway in early second language acquisition. Communications Biology, 7(1), 1298.

38. Grant, A. M., Kousaie, S., Coulter, K., Gilbert, A. C., Baum, S. R., Gracco, V., Titone, D., Klein, D., & Phillips, N. A. (2022). Age of Acquisition Modulates Alpha Power During Bilingual Speech Comprehension in Noise. Frontiers in Psychology, 13, 865857.

39. Green, D. W., & Abutalebi, J. (2013). Language control in bilinguals: The adaptive control hypothesis. Journal of Cognitive Psychology, 25(5), 515–530.

40. Gullifer, J. W., Chai, X. J., Whitford, V., Pivneva, I., Baum, S., Klein, D., & Titone, D. (2018). Bilingual experience and resting-state brain connectivity: Impacts of L2 age of acquisition and social diversity of language use on control networks. Neuropsychologia, 117, 123–134.

41. Harmony, T. (2013). The functional significance of delta oscillations in cognitive processing. Frontiers in Integrative Neuroscience, 7.

42. Har-shai Yahav, P., & Zion Golumbic, E. (2021). Linguistic processing of task-irrelevant speech at a cocktail party. eLife, 10, e65096.

43. Hartanto, A., & Yang, H. (2016). Disparate bilingual experiences modulate task-switching advantages: A diffusion-model analysis of the effects of interactional context on switch costs. Cognition, 150, 10–19.

44. Hauswald, A., Keitel, A., Chen, Y., Rösch, S., & Weisz, N. (2022). Degradation levels of continuous speech affect neural speech tracking and alpha power differently. European Journal of Neuroscience, 55(11–12), 3288–3302.

45. Hosoda, C., Tanaka, K., Nariai, T., Honda, M., & Hanakawa, T. (2013). Dynamic Neural Network Reorganization Associated with Second Language Vocabulary Acquisition: A Multimodal Imaging Study. The Journal of Neuroscience, 33(34), 13663–13672.

46. Kaushik, P., Moye, A., Vugt, M. V., & Roy, P. P. (2022). Decoding the cognitive states of attention and distraction in a real-life setting using EEG. Scientific Reports, 12(1), 20649.

47. Kheder, S., & Kaan, E. (2021). Cognitive control in bilinguals: Proficiency and code-switching both matter. Cognition, 209, 104575.

48. Klimesch, W. (1999). EEG alpha and theta oscillations reflect cognitive and memory performance: A review and analysis. Brain Research Reviews, 29(2–3), 169–195.

49. Klimesch, W. (2012). Alpha-band oscillations, attention, and controlled access to stored information. Trends in Cognitive Sciences, 16(12), 606–617.

50. Klimovich-Gray, A., Barrena, A., Agirre, E., & Molinaro, N. (2021). One Way or Another: Cortical Language Areas Flexibly Adapt Processing Strategies to Perceptual And Contextual Properties of Speech. Cerebral Cortex, 31(9), 4092–4103.

51. Knyazev, G. G. (2012). EEG delta oscillations as a correlate of basic homeostatic and motivational processes. Neuroscience & Biobehavioral Reviews, 36(1), 677–695.

52. Korenar, M., Treffers-Daller, J., & Pliatsikas, C. (2023). Dynamic effects of bilingualism on brain structure map onto general principles of experience-based neuroplasticity. Scientific Reports, 13(1), 3428.

53. Kösem, A., Basirat, A., Azizi, L., & Van Wassenhove, V. (2016). High-frequency neural activity predicts word parsing in ambiguous speech streams. Journal of Neurophysiology, 116(6), 2497–2512.

54. Krause, C. M., Sillanmäki, L., Koivisto, M., Saarela, C., Häggqvist, A., Laine, M., & Hämäläinen, H. (2000). The effects of memory load on event-related EEG desynchronization and synchronization. Clinical Neurophysiology, 111(11), 2071–2078.

55. Krizman, J., Marian, V., Shook, A., Skoe, E., & Kraus, N. (2012). Subcortical encoding of sound is enhanced in bilinguals and relates to executive function advantages. Proceedings of the National Academy of Sciences, 109(20), 7877–7881.

56. Krizman, J., Skoe, E., Marian, V., & Kraus, N. (2014). Bilingualism increases neural response consistency and attentional control: Evidence for sensory and cognitive coupling. Brain and Language, 128(1), 34–40.

57. Krizman, J., Tierney, A., Nicol, T., & Kraus, N. (2021). Listening in the Moment: How Bilingualism Interacts With Task Demands to Shape Active Listening. Frontiers in Neuroscience, 15, 717572.

58. Kuipers, J. R., & Thierry, G. (2015). Bilingualism and increased attention to speech: Evidence from event-related potentials. Brain and Language, 149, 27–32.

59. Lalor, E. C., & Foxe, J. J. (2010). Neural responses to uninterrupted natural speech can be extracted with precise temporal resolution. European Journal of Neuroscience, 31(1), 189–193.

60. Lenth, R., Singmann, H., Love, J., Buerkner, P., & Herve, M. (2019). Emmeans: estimated marginal means, aka least-squares means (Version 1.3. 4). Emmeans Estim Marg Means Aka Least-Sq Means, 4.

61. Li, P., Legault, J., & Litcofsky, K. A. (2014). Neuroplasticity as a function of second language learning: Anatomical changes in the human brain. Cortex, 58, 301–324.

62. Liu, C., Jiao, L., Li, Z., Timmer, K., & Wang, R. (2021). Language control network adapts to second language learning: A longitudinal rs-fMRI study. Neuropsychologia, 150, 107688.

63. Lizarazu, M., Carreiras, M., Bourguignon, M., Zarraga, A., & Molinaro, N. (2021). Language Proficiency Entails Tuning Cortical Activity to Second Language Speech. Cerebral Cortex, 31(8), 3820–3831.

64. Lu, L., Deng, Y., Xiao, Z., Jiang, R., & Gao, J.-H. (2023). Neural Signatures of Hierarchical Linguistic Structures in Second Language Listening Comprehension. Eneuro, 10(6), ENEURO.0346-22.2023.

65. Luk, G., Anderson, J. A. E., Craik, F. I. M., Grady, C., & Bialystok, E. (2010). Distinct neural correlates for two types of inhibition in bilinguals: Response inhibition versus interference suppression. Brain and Cognition, 74(3), 347–357.

66. Luk, G., Bialystok, E., Craik, F. I. M., & Grady, C. L. (2011). Lifelong Bilingualism Maintains White Matter Integrity in Older Adults. The Journal of Neuroscience, 31(46), 16808–16813.

67. Mai, G., Minett, J. W., & Wang, W. S.-Y. (2016). Delta, theta, beta, and gamma brain oscillations index levels of auditory sentence processing. NeuroImage, 133, 516–528.

68. Mai, G., & Wang, W. S. -Y. (2023). Distinct roles of delta- and theta-band neural tracking for sharpening and predictive coding of multi-level speech features during spoken language processing. Human Brain Mapping, 44(17), 6149–6172.

69. Mai, G., & Wang, W. S. (2019). Delta and theta neural entrainment during phonological and semantic processing in speech perception. BioRxiv, 556837.

70. Mårtensson, J., Eriksson, J., Bodammer, N. C., Lindgren, M., Johansson, M., Nyberg, L., & Lövdén, M. (2012). Growth of language-related brain areas after foreign language learning. NeuroImage, 63(1), 240–244.

71. Mechelli, A., Crinion, J. T., Noppeney, U., O’Doherty, J., Ashburner, J., Frackowiak, R. S., & Price, C. J. (2004). Structural plasticity in the bilingual brain. Nature, 431(7010), 757–757.

72. Meyer, L. (2018). The neural oscillations of speech processing and language comprehension: State of the art and emerging mechanisms. European Journal of Neuroscience, 48(7), 2609–2621.

73. Molinaro, N., & Lizarazu, M. (2018). Delta(but not theta)-band cortical entrainment involves speech-specific processing. European Journal of Neuroscience, 48(7), 2642–2650.

74. Mouthon, M., Khateb, A., Lazeyras, F., Pegna, A. J., Lee-Jahnke, H., Lehr, C., & Annoni, J. M. (2020). Second-language proficiency modulates the brain language control network in bilingual translators: an event-related fMRI study. Bilingualism: Language and cognition, 23(2), 251–264.

75. Obleser, J., Wöstmann, M., Hellbernd, N., Wilsch, A., & Maess, B. (2012). Adverse Listening Conditions and Memory Load Drive a Common Alpha Oscillatory Network. The Journal of Neuroscience, 32(36), 12376–12383.

76. Olguin, A., Bekinschtein, T. A., & Bozic, M. (2018). Neural Encoding of Attended Continuous Speech under Different Types of Interference. Journal of Cognitive Neuroscience, 30(11), 1606–1619.

77. Olguin, A., Cekic, M., Bekinschtein, T. A., Katsos, N., & Bozic, M. (2019). Bilingualism and language similarity modify the neural mechanisms of selective attention. Scientific Reports, 9(1), 8204.

78. Olsen, R. K., Pangelinan, M. M., Bogulski, C., Chakravarty, M. M., Luk, G., Grady, C. L., & Bialystok, E. (2015). The effect of lifelong bilingualism on regional grey and white matter volume. Brain Research, 1612, 128–139.

79. Oostenveld, R., Fries, P., Maris, E., & Schoffelen, J. M. (2011). FieldTrip: open source software for advanced analysis of MEG, EEG, and invasive electrophysiological data. Computational intelligence and neuroscience, 2011(1), 156869.

80. O’Sullivan, J. A., Power, A. J., Mesgarani, N., Rajaram, S., Foxe, J. J., Shinn-Cunningham, B. G., Slaney, M., Shamma, S. A., & Lalor, E. C. (2015). Attentional Selection in a Cocktail Party Environment Can Be Decoded from Single-Trial EEG. Cerebral Cortex, 25(7), 1697–1706.

81. Peelle, J. E., Gross, J., & Davis, M. H. (2013). Phase-Locked Responses to Speech in Human Auditory Cortex are Enhanced During Comprehension. Cerebral Cortex, 23(6), 1378–1387.

82. Perani, D., & Abutalebi, J. (2015). Bilingualism, dementia, cognitive and neural reserve. Current Opinion in Neurology, 28(6), 618–625.

83. Pfurtscheller, G., & Da Silva, F. L. (1999). Event-related EEG/MEG synchronization and desynchronization: basic principles. Clinical neurophysiology, 110(11), 1842–1857.

84. Phelps, J., Attaheri, A., & Bozic, M. (2022). How bilingualism modulates selective attention in children. Scientific Reports, 12(1), 6381.

85. Phelps, J., & Bozic, M. (2025). Flexible functional adaptation of selective attention in bilingualism. Bilingualism: Language and Cognition, 28(2), 312–326.

86. Pliatsikas, C. (2020). Understanding structural plasticity in the bilingual brain: The Dynamic Restructuring Model. Bilingualism: Language and Cognition, 23(2), 459–471.

87. Pliatsikas, C., DeLuca, V., Moschopoulou, E., & Saddy, J. D. (2017). Immersive bilingualism reshapes the core of the brain. Brain Structure and Function, 222(4), 1785–1795.

88. Pliatsikas, C., Moschopoulou, E., & Saddy, J. D. (2015). The effects of bilingualism on the white matter structure of the brain. Proceedings of the National Academy of Sciences, 112(5), 1334–1337.

89. Pion-Tonachini, L., Kreutz-Delgado, K., & Makeig, S. (2019). ICLabel: An automated electroencephalographic independent component classifier, dataset, and website. NeuroImage, 198, 181–197.

90. Rimmele, J. M., Golumbic, E. Z., Schröger, E., & Poeppel, D. (2015). The effects of selective attention and speech acoustics on neural speech-tracking in a multi-talker scene. Cortex, 68, 144–154.

91. Rossi, E., Pereira Soares, S. M., Prystauka, Y., Nakamura, M., & Rothman, J. (2023). Riding the (brain) waves! Using neural oscillations to inform bilingualism research. Bilingualism: Language and Cognition, 26(1), 202–215.

92. Schlegel, A. A., Rudelson, J. J., & Tse, P. U. (2012). White Matter Structure Changes as Adults Learn a Second Language. Journal of Cognitive Neuroscience, 24(8), 1664–1670.

93. Stein, M., Federspiel, A., Koenig, T., Wirth, M., Strik, W., Wiest, R., … & Dierks, T. (2012). Structural plasticity in the language system related to increased second language proficiency. Cortex, 48(4), 458–465.

94. StrauÃŸ, A., WÃ¶stmann, M., & Obleser, J. (2014). Cortical alpha oscillations as a tool for auditory selective inhibition. Frontiers in Human Neuroscience, 8.

95. Sulpizio, S., Del Maschio, N., Del Mauro, G., Fedeli, D., & Abutalebi, J. (2020). Bilingualism as a gradient measure modulates functional connectivity of language and control networks. NeuroImage, 205, 116306.

96. Tao, L., Wang, G., Zhu, M., & Cai, Q. (2021). Bilingualism and domain-general cognitive functions from a neural perspective: A systematic review. Neuroscience & Biobehavioral Reviews, 125, 264–295.

97. Tu, L., Wang, J., Abutalebi, J., Jiang, B., Pan, X., Li, M., Gao, W., Yang, Y., Liang, B., Lu, Z., & Huang, R. (2015). Language exposure induced neuroplasticity in the bilingual brain: A follow-up fMRI study. Cortex, 64, 8–19.

98. Tune, S., Alavash, M., Fiedler, L., & Obleser, J. (2021). Neural attentional-filter mechanisms of listening success in middle-aged and older individuals. Nature Communications, 12(1), 4533.

99. Wei, X., Gunter, T. C., Adamson, H., Schwendemann, M., Friederici, A. D., Goucha, T., & Anwander, A. (2024). White matter plasticity during second language learning within and across hemispheres. Proceedings of the National Academy of Sciences, 121(2), e2306286121.

100. Wisniewski, M. G., Thompson, E. R., & Iyer, N. (2017). Theta- and alpha-power enhancements in the electroencephalogram as an auditory delayed match-to-sample task becomes impossibly difficult. Psychophysiology, 54(12), 1916–1928.

101. Wöstmann, M., Lim, S.-J., & Obleser, J. (2017). The Human Neural Alpha Response to Speech is a Proxy of Attentional Control. Cerebral Cortex, 27(6), 3307–3317.

